# ALS monocyte-derived microglia reveal cytoplasmic TDP-43 accumulation, DNA damage, and cell-specific impairment of phagocytosis associated with disease progression

**DOI:** 10.1101/2020.10.25.354399

**Authors:** Hazel Quek, Carla Cuní-López, Romal Stewart, Tiziana Colletti, Antonietta Notaro, Yifan Sun, Christine C. Guo, Michelle K. Lupton, Tam Hong Nguyen, Lotta E. Oikari, Tara L. Roberts, Yi Chieh Lim, Vincenzo La Bella, Anthony R. White

## Abstract

**Aims:** Amyotrophic lateral sclerosis (ALS) is a multifactorial neurodegenerative disease characterised by the loss of upper and lower motor neurons. Neuroinflammation mediated by microglial activation is evident in post-mortem brain tissues, and in brain imaging of patients with ALS. However, the exact role of microglia in ALS remains to be elucidated partly due to the lack of an accurate microglial model system that is able to recapitulate the clinical pathology of ALS. Moreover, direct sampling of microglia from patients with ALS is not feasible, further limiting the study of microglial function in ALS. To address this shortcoming, we describe an approach that generates monocyte-derived microglia (MDMi) that are capable of expressing molecular markers, and functional characteristics similar to resident human brain microglia. Importantly, MDMi can be routinely and reproducibly generated from ALS patient blood, and reveal patient heterogeneity associated with age, sex and disease subgroup.

**Methods:** MDMi were successfully established from all 30 ALS patients, including 15 patients with slow disease progression, 6 with intermediate progression, and 9 with rapid progression, together with 20 non-affected heathy controls (HC).

**Results:** Our ALS MDMi model recapitulated canonical pathological features of ALS including non-phosphorylated and phosphorylated-TDP-43-positive pathological inclusions. We further observed significantly impaired phagocytosis, altered cytokine expression and microglial morphology, as well as elevated DNA damage in ALS compared to HC MDMi. Abnormal phagocytosis was observed in all ALS cases, and was correlated to the progression of disease. Moreover, in-depth analysis of individual microglia revealed cell-specific variation in phagocytic function that was significantly altered, and exacerbated in rapid disease progression.

**Conclusions:** Our approach enabled us to generate ALS patient microglia from peripheral blood samples using a rapid, robust, cost-effective, and reproducible protocol. We have shown that ALS monocyte-derived microglia have significantly altered functional behaviour compared to age-matched HCs, with a major deficit in phagocytic activity. This is also the first demonstration of abnormal TDP-43 localisation in microglia grown from ALS patients. Overall, this approach is highly applicable to monitor disease progression and can be applied as a functional readout in clinical trials for anti-neuroinflammatory agents. Additionally, this model system can be used as a basis for personalised therapeutic treatment for ALS, as well as other neurodegenerative diseases.

## Introduction

Amyotrophic lateral sclerosis (ALS) is a debilitating disease characterised by the loss of motor neurons in the brain and spinal cord, resulting in progressive muscle weakness and eventual death. Neuroinflammation, a key contributor to the pathological changes in ALS is mediated by microglia, the resident innate immune cells of the central nervous system (CNS). Under normal conditions, these cells survey and maintain the brain microenvironment by pruning neuronal synapses, phagocytosing cellular debris, and responding to signalling molecules to maintain cellular homeostasis. However, during neuroinflammation, microglia respond to activating signals, exhibit an altered morphology, and mediate inflammatory responses by secreting pro- and/or anti-inflammatory cytokines in order to restore brain homeostasis (Ginhoux et al., 2010). Moreover, ALS pathology encompasses a complex process involving both neuronal and non-neuronal changes in the brain and spinal cord of affected individuals. The non-neuronal contribution to the disease progression of ALS is supported by the infiltration of myeloid cells from circulation into the CNS, as well as microgliosis, a neuroinflammatory process resulting in the increased numbers of activated microglia. There is increasing evidence that these changes alter disease progression and severity (Beers and Appel 2019; Henkel et al. 2004). However, it remains difficult to delineate exactly how microglia contribute to the progression of pathological changes observed in ALS.

Until recently, research carried out to investigate the involvement of microglia in ALS relied predominantly on animal models, and post-mortem human CNS tissue. Sampling of microglia from human brain autopsy and biopsy tissue is not practical for high-throughput screening platforms, while culturing isolated human microglia ex *vivo* is challenging due to its restricted proliferative capacity, cell viability, and rapid changes to its unique CNS identity once removed from the brain microenvironment (O. Butovsky et al., 2012). New approaches using induced pluripotent stem cell (iPSC)-derived microglia are now available but have yet to provide major insights into ALS neuroinflammatory processes, likely due to the relative complexity, high variability and increased time-frames required to generate iPSC-derived microglia. Moreover, iPSC-derived microglia may not accurately recapitulate the heterogeneity of clinical features observed in this disease due to the loss of epigenetic factors during reprogramming from skin cells to stem cells (Hawrot, Imhof, & Wainger, 2020; Philips & Rothstein, 2015). Hence, in an attempt to address these short-comings and provide further insights into the pathological role of microglia in ALS, we generated a patient-derived microglia model using peripheral blood-derived monocytes. This monocyte-derived microglia (MDMi) model is a rapid, minimally-invasive system that allows for multiple sampling at various stages of ALS progression. This in turn would deliver better representation of changes that may occur in microglia during the progression of disease thereby providing better clinical outcomes.

We show that MDMi express key features of ALS pathology including microglia activation, as demonstrated by morphological changes, together with altered cytokine expression. Importantly, we were able to recapitulate the aberrant accumulation of cytoplasmic TAR DNA binding protein of 43 kDA (TDP-43), and phosphorylated TDP-43 inclusions in ALS MDMi, a key pathological hallmark present in the majority of ALS cases. In addition, ALS MDMi also showed significantly impaired phagocytosis, DNA damage and inflammasome activation. These findings demonstrate that the MDMi provide a novel platform for recapitulating several major pathological features observed in ALS and provide a novel platform to delineate the functional role of microglia in ALS and other neurological disorders.

## Materials and methods

### Patient recruitment

This study involved the recruitment of an ALS patient cohort from the ALS clinical Research Centre in Palermo, Italy. Healthy control (HC) participants with no clinical symptoms were recruited from either the ALS Clinical Research Centre, or the *Prospective Imaging Studying of Aging: Genes, Brain and Behaviour* study (PISA) at QIMR Berghofer Medical Research Institute, Queensland, Australia. All research adhered to the ethical guidelines on human research outlined by the National Health and Medical Research Council of Australia (NHMRC). Ethical approval was obtained from QIMR Berghofer Medical Research Institute, and University of Palermo. All participants provided informed consent before participating in the study.

The methodology for PISA recruitment has been described previously (Lupton et al., 2020). For ALS patients, the degree of functional impairment was assessed with the Revised Amyotrophic Lateral Sclerosis Functional Rating Scale (ALSFRS-R) (Cedarbaum et al., 1999). The rate of disease progression was evaluated using the ALSFRS-R score: progression rate ratio (Delta FS, ΔFS) according to the following formula: (48-ALSFRS-R score at time of diagnosis)/ time onset to diagnosis). Patients were then stratified into three arbitrary groups according to ΔFS: slow (ΔFS < 0.5), intermediate (ΔFS= 0.5-1.0), rapid (ΔFS >1.0), which can predict life expectancy post diagnosis (Kimura et al., 2006; Kollewe et al., 2008).

A summary of disease characteristics, and demographics of the patients and control donors used in this study, are shown in Table 1. HCs were defined as donors without clinical symptoms as confirmed by retrospective medical review from each study cohort.

**Table 1.** Summary of donor information

| **Study Cohorts:** | **Healthy Control (HC)** | **ALS** |  |  |
| --- | --- | --- | --- | --- |
| **Disease progression** |  | **Slow** | **Intermediate** | **Rapid** |
| **No. of participants** | **N=20** | **N=15** | **N=6** | **N=9** |
| **Sex of participants** |  |  |  |  |
| **Females** | 50% (11/20) | 33.3% | 66.7% | 44.4% |
| **%, (proportion)** |  | (5/15) | (4/6) | (4/9) |
| **Males** | 45% (9/20) | 66.7% | 33.3% | 55.6% |
| **%, (proportion)** |  | (10/15) | (2/6) | (5/9) |
| **Age: Mean ( ± SD)** | 65.3 (8.9) | 57.2 (10.2) | 70.3 (9.7) | 66.0 (8.84) |
| **Site of onset** |  |  |  |  |
| **Bulbar: %, (proportion)** |  | 13.3% (2) | 16.7% (1) | 22.2% (2) |
| **Spinal: %, (proportion)** |  | 86.7% (13) | 83.3% (5) | 77.8% (7) |
| **ALSFRS-R<sup>a</sup>: mean, ± SD (range)** | | 41.6, ±3.4 (34-46) | 41, ±2.3 (38-44) | 36.8, ±5.04 (30-46) |
| **ΔFS<sup>b</sup>: mean, ± SD (range)** | | 0.25, ±0.1 (0.1-0.45) | 0.60, ± 0.1 (0.57-0.67) | 1.72, 0.76 (1-3.4) |

<sup>a</sup>ALSFRS-R (at diagnosis, when PBMCs were taken; normal score=48)
<sup>b</sup>Delta FS ratio (rate of disease progression); (48- ALSFRS-R score at time of diagnosis)/ time onset to diagnosis)
SD, Standard deviation

### Isolation of PBMCs from blood samples

Donor blood was collected into tubes containing ethylenediaminetetraacetic acid (EDTA) (Becton-Dickson, NJ, USA) by a qualified phlebotomist. Briefly, donor blood was diluted 1: 1 with phosphate buffered saline (PBS) and transferred into sterile SepMate 50 (STEMCELL Technologies, BC, Canada) as per manufacturer’s instructions. Peripheral blood monocytes (PBMCs) were collected and resuspended in freezing media (90% v/v fetal bovine serum (FBS, Thermofisher, CA, USA) supplemented with 10% dimethyl sulfoxide (DMSO, Merck KGaA, Hesse, Germany) and cryopreserved in liquid nitrogen.

### Generation of Monocyte-Derived Microglia (MDMi)

Briefly, cyropreserved PBMCs were thawed at 37°C and diluted into culture media and centrifugated at 300g for 5 min. Supernatant was aspirated completely and pellet was resuspended in culture media. Cells were plated onto Matrigel™ coated (Corning, NY, USA) 48-well plates and incubated in cell culture incubator of 37°C, 5% CO2. For microglia differentiation, culture media supplemented with 0.1μg/ml of inter-leukin (IL)-34 (Lonza, Basel-Stadt, Switzerland) and 0.01μg/ml of granulocyte-macrophage colony-stimulating factor (GM-CSF) (Lonza, Basel-Stadt, Switzerland) was then added to these cells. All cells were cultured under standard culture conditions for up to 14 days. Differentiation of PBMCs into MDMi was confirmed by 1) the expression of classical ramified morphology, 2) immunofluorescence for microglial markers, and 3) mRNA expression of microglial specific genes in comparison to monocyte-derived macrophages (MDMa) and monocyte-derived dendritic cells (MDDCs).

All samples used for downstream assays were grouped according to age, sex, and disease subgroup (slow, intermediate, and rapid) with details listed in the respective figure legends.

### Generation of monocyte-derived macrophages (MDMa) and dendritic cells (MDDCs)

For MDMa induction, PBMCs were seeded in a 48-well plate with culture media. 10ng/ml GM-CSF was added into the culture medium and was further differentiated for 14 days. MDMa have a classic large cell surface area and amoeboid morphology that are distinct from MDMi.

For MDDC induction, PBMCs were seeded in a 6-well plate with culture media. 50ng/ml of IL-4, and 100ng/ml of GM-CSF were added, and cultured until day 7, after which cells were either harvested for downstream assays or induced with 100ng/ml LPS and 1000IU/ml IFNγ for 48 hr for MDDC maturation. Mature MDDCs showed characteristic stellate morphology (multiple pseudopodia), distinct from MDMa and MDMi.

### TDP-43 treatment

Healthy MDMi were differentiated until day 14. Human recombinant TDP-43/TARDBP protein (R&D systems, MN, US) was reconstituted as per manufacturer’s instruction. Briefly, TDP-43 recombinant protein utilised at 1nM and 10nM was added into MDMi cultures for a period of 24 hr.

### Immunofluorescence

Cells were plated onto 8-well chamber slides (Ibidi, DKSH, Germany), and immunofluorescence was performed as previously described (Quek et al., 2017). Briefly, cells were fixed with either ice-cold methanol or 4% paraformaldehyde in PBS. Permeabilisation of PFA fixed samples was performed with PBS containing 0.3% Triton-X 100. Samples were blocked with PBS containing 5% bovine serum albumin (BSA) (Sigma, USA). Primary antibodies used were as follows: Anti-P2RY12 (Alomone Labs, #APR-20, 1:200), Anti-IBA1 (Abcam, #Ab5076, 1:500; Wako, #019-19741, 1:500), Anti-TDP-43 (Cosmo Bio, #TIP-TD-P09, 1/500), Anti-p-TDP43 (Cosmo Bio, #TIP-TD-P09, 1:500), Anti-γH2AX (EMD Millipore, #05-636, 1:400), Anti-ASC (AdipoGen, #AG-25B-0006-C100, 1:200), Anti-NLRP3 (AdipoGen, #AG-20B-0014-C100, 1:500) were incubated overnight. Cells were washed thrice with 0.1% Triton-X in PBS followed by incubation with secondary antibodies and DAPI (nuclear dye) (Sigma, USA) for 2 hr at room temperature. Antibody specificity was confirmed by performing secondary antibody only controls. Investigators were blinded to the conditions of the experiments during data collection and analyses. Fluorescence images were captured with a confocal laser scanning microscope (LSM-780, Carl Zeiss). All settings were kept consistent during acquisition.

### Live cell imaging of phagocytosis

Characterisation of phagocytic function was examined using the live cell imaging on the IncuCyte ZOOM (Essen Biosciences). Briefly, sonicated fluorescent pHrodo-labelled *E.coli* particles (Life Technologies, CA, USA) were added to MDMi cell cultures. Multiple images were captured every hour at 10x magnification for a total of 15 hr using standard phase contrast and red fluorescence settings. Data were analysed using the IncuCyte ZOOM software 2018A. Background fluorescence was subtracted from original images using the top-Hat threshold transform algorithm. All parameters were kept constant between different experiments. For each time point, normalised phagocytosis was calculated as (Red object area)/ (Phase object count).

### RNA extraction and quantitative real-time polymerase chain reaction (qRT-PCR)

RNA extraction was performed according to (Quek et al, 2016). Briefly, total RNA extraction was performed using Direct-zol RNA miniprep kit (Integrated Sciences, Australia) according to the manufacturer’s instructions. cDNA synthesis was then performed using a SensiFast cDNA synthesis kit (Bioline, London, UK). Samples were diluted and mixed with SensiFAST Sybr Lo-Rox master mix before loading as triplicates for qRT-PCR. qRT-PCR was performed using Applied Biosystems ViiA 7 system. Similar *18S* ribosomal RNA expression was observed in HC and in ALS MDMi, hence was used as a housekeeping gene. ΔΔCT values normalised to *18S* were used to assess relative gene expression between samples. Samples with cycle threshold (CT) values more than two times standard deviation of the average of each reference gene were excluded from analysis. Melt curve analyses confirmed a single melt curve peak for all primers, summarised in Supplementary Table 1.

### Skeleton analysis

Phase contrast images of MDMi were captured using a spinning disc confocal microscope using a 20x objective of 0.4 numerical aperture. The characterisation of microglia morphology was performed using ImageJ analysis software (ImageJ 1.52n, National Institutes of Health, Maryland, USA). Phase contrast images of MDMi were first pre-processed using a macro script that applied a threshold, followed by conversion to binary images. The “AnalyzeSkeleton” plugin was then executed on the binarised images to analyse MDMi branch length, branch number (cell process and end-points per cell), and branch junctions (triple or quadruple junctions). These data measure microglial morphology (complexity and process length) (Young & Morrison, 2018). One hundred cells per patient or individual in each subgroup were analysed.

### Statistical Analysis

Statistical analysis of the differences between groups was performed using the Student’s t-test (two-tailed), or the Mann-Whitney U when appropriate. One-way analysis of variance (ANOVA) followed by Dunnett’s multiple comparison test was used to compare multiple treatment groups versus control. Normal distribution of qPCR data after log-transformation was assumed. Corresponding statistical test/s and accompanying significance data are listed in each figure legend. All statistical analyses were performed using GraphPad Prism 8 (Graphpad Software). *\*P* <0.05 was considered statistically significant. All data were presented as mean ± SEM.

## Results

### Generation of human monocyte-derived microglia (MDMi)

Initial generation and characterisation of MDMi was performed with PBMCs collected from healthy volunteers. Isolated PBMCs were cultured for up to 14 days with induction media supplemented with 100ng/ml IL-34 and 10ng/ml GM-CSF to generate a microglia-like phenotype, which include the presence of a small soma with ramified morphology compared to day 7 MDMi (early stage differentiation) (Fig. 1a), and positive immunostaining for markers enriched in microglia such as, P2ry12 and Iba1 (Fig. 1b). Expression levels of seminal microglial markers including *PROS1, GPR34, C1QA, MERTK, GAS6, APOE* and *P2RY12* (Oleg Butovsky et al., 2014; Ormel et al., 2020; Ryan et al., 2017; Sellgren et al., 2017), and well-known microglial genes such as *TREM2, CD68,* and *HLA-DRA* were upregulated in MDMi at day 14 compared to isolated monocytes. Additionally, we evaluated microglial transcriptional genes such as *RUNX1, PU.1* and *IRF8* in day 14 MDMi compared to isolated monocytes (Ginhoux et al., 2010; Wehrspaun, Haerty, & Ponting, 2015) (Fig. 1c, S1a). *RUNX1*, a key regulator of myeloid proliferation and differentiation was downregulated in MDMi compared to monocytes, indicating a mature and resting microglial phenotype (Zusso et al., 2012). There were no changes in *PU.1* and *IRF8* expression levels between monocytes and MDMi indicating that both transcription factors are important in regulating monocytes and microglia (Kierdorf et al., 2013). Overall, these results indicate that MDMi display a mature microglial phenotype.

**Figure 1.**
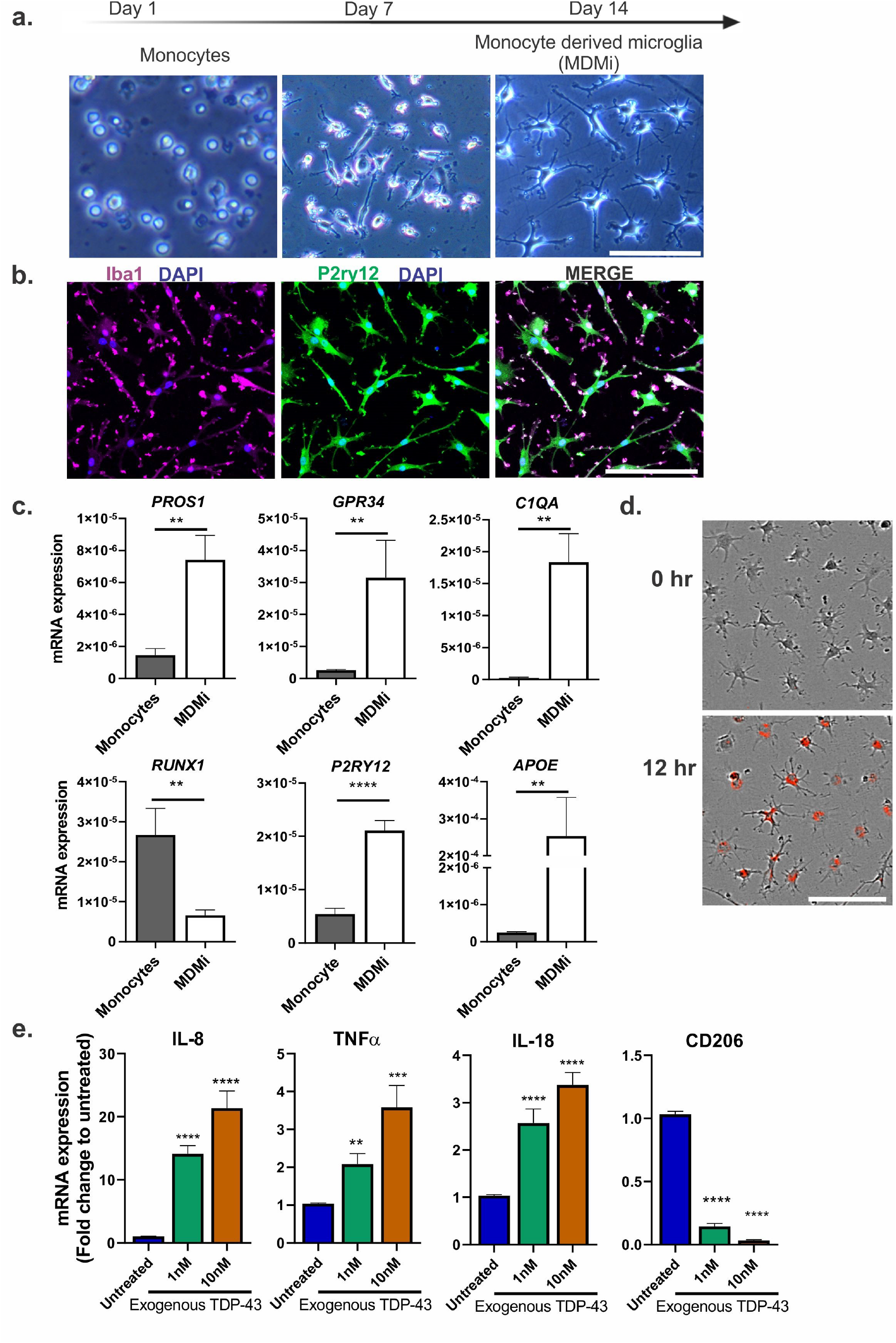
Generation and characterisation of human monocyte-derived microglia (MDMi). **(a)** Schematic timeline of MDMi differentiation for 1, 7 and 14 days in culture, with representative phase contrast images. **(b)** Immunofluorescence images of Iba1, P2ry12 with counter-stain DAPI in MDMi (MDMi from <40 years of age: n=5). **(c)** Gene expression of seminal microglia genes including *PROS1, GPR34, C1QA, RUNX1, P2RY12* and *APOE* between isolated monocytes and MDMi (isolated monocytes and MDMi from <40 years of age: n=5). **(d)** Representative phase contrast images of MDMi with pHrodo-labelled *E.coli* particles (red) uptake at 12 hr after treatment compared to untreated (top) (MDMi from <40 years of age: n=5). **(e)** Gene expression of MDMi treated with 1nM (green) and 10nM (orange) TDP-43 recombinant protein (<40 years of age; n=3). The y-axis represents the fold change of mRNA expression levels (*IL-8, TNFα, IL-18* and *CD206)* normalised to untreated cells over 24 hr treatment. Statistical analysis between two groups was performed using Student’s *t* test and between multiple groups using one-way ANOVA. Values are mean ± SEM (*P < 0.05, ** *P* < 0.01, *** *P* < 0.001, **** *P* < 0.0001). Scale bars= 50μm.

As monocytes can also commit to different tissue-specific macrophage lineages influenced by cytokine mediators, we further compared MDMi with monocyte-derived macrophages (MDMa), and monocyte-derived dendritic cells (MDDCs) (Fig. S1b). We show that MDMa display decreased expression of microglia-markers, *TREM2, CX3CR1* and *CD68,* while increased expression of macrophage-marker, *CD45* compared to MDMi (Fig. S1c). Additionally, we show that MDMi express both *CD209,* an immature MDDC marker and *CCR7,* a mature MDDC marker at very low levels, but expressed *CD68,* a microglial marker at a higher level compared to MDDCs (Fig. S1d), confirming the specificity of our MDMi population that is different from MDMa and MDDCs. Overall, these results show that MDMi display characteristic microglial genes expressed at a higher level compared to other cells within the monocyte lineage.

Key functions of microglia include phagocytosis and release of inflammatory cytokines in response to stimuli. To validate the functional capacity of MDMi cells, we stimulated MDMi with pHrodo-labelled *E.coli* particles, where uptake into the acidic phagosomes results in fluorescence. An increased in red intensity over time confirmed the phagocytic ability of MDMi (Fig 1d). Further, to examine response of MDMi to various disease-specific damage-associated molecular patterns (DAMPs), we treated MDMi with TDP-43 recombinant protein at 1 and 10nM. This showed a dose-dependent increase of pro-inflammatory cytokines (IL-8, TNFα and IL-18) and decrease in anti-inflammatory cell surface marker (CD206) when compared to untreated MDMi. (Fig. 1e). Taken together, these results confirmed that MDMi have a similar functional capacity to brain microglia.

### ALS patient-derived MDMi reveal morphological differences compared to HCs

As they mature, microglia lose their ability to proliferate (Bennett et al., 2016). Hence, to determine the proliferative capacity of MDMi from ALS patients and HCs, we examined Ki67 expression, an indicator of cell proliferation. Similar expression levels of Ki67 was observed across HC and ALS disease subgroups, confirming the presence of mature differentiated MDMi in all cases (Fig S2a).

Changes in microglial morphology have been shown to reflect the health of the brain (Frank-Cannon, Alto, McAlpine, & Tansey, 2009). In response to damage or an injury, microglia undergo a rapid transition from a ramified to an amoeboid form, characteristic of its activated state (Nimmerjahn, Kirchhoff, & Helmchen, 2005). To examine microglial morphology, ALS MDMi were stratified into three subgroups with different rates of disease progression (rapid, intermediate and slow) (Table 1, Fig. 2a, S2b). We observed a significant reduction in microglial branch length (P <0.0001), and end-points (P <0.0001) in ALS cultures when compared to age-matched HCs (Fig.2b-c, S2c-e). This was also associated with a decrease in the number of microglial branches per cell (P <0.0001) in ALS compared to HC MDMi, suggesting a disease-specific morphological change in ALS MDMi (Fig. 2d, S2f). Notably, a portion of ALS MDMi within each subgroup presented a similar microglial morphology to HC MDMi, demonstrating heterogeneity of cell morphologies within each disease subgroup (Fig. S2d-e).

**Figure 2.**
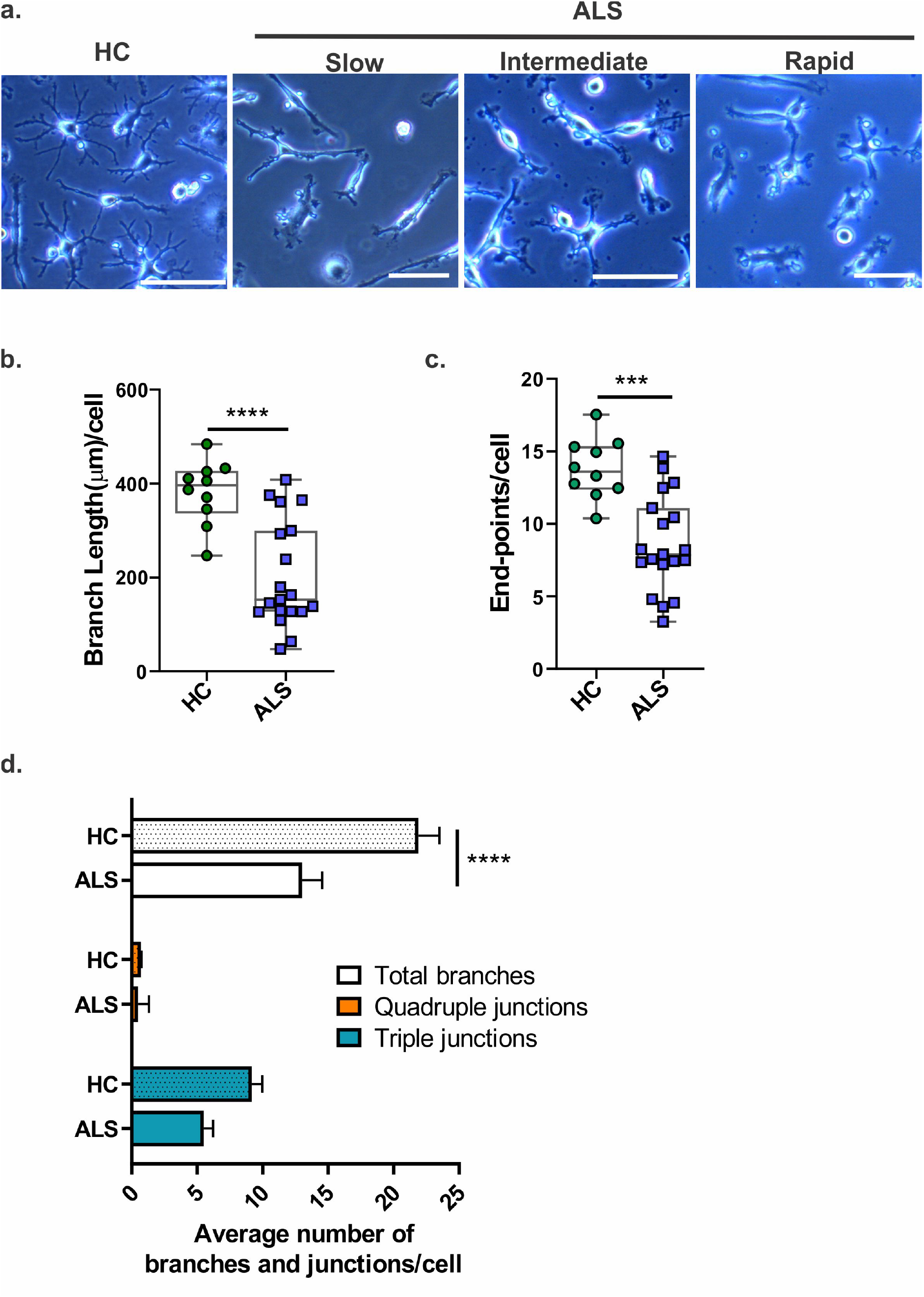
Morphology of ALS MDMi. **(a)** Phase contrast images of mature ALS MDMi subgroups (slow, intermediate, and rapid) 14 days in culture. **(b)** Microglial branch length of HC and ALS MDMi were analysed by ImageJ and normalised to total length per cell; HC: n=10, ALS: n=19. **(c)** Microglial end-points of ALS and HC MDMi were analysed by ImageJ and normalised against cell number; HC: n=10, ALS: n=19. **(d)** Average number of branches, and secondary junctions of HC and ALS MDMi were analysed by ImageJ and normalised against cell number; HC: n=10, ALS: n=19. Statistical analysis between two groups was performed using Student’s *t* test and between multiple groups using one-way ANOVA. Values are mean ± SEM (***P < 0.001, **** *P* < 0.0001). Scale bars= 50μm.

In addition, we observed a significant reduction in microglial branch length and end-points in ALS across all three disease subgroups compared to HC MDMi (Fig. S2d-f). However, no subgroup specific trends were observed (Fig. S2d-f). Interestingly, we observed a significant reduction in microglial branch junctions (triple junctions (P=0.0008), and quadruple junctions (P=0.0002)) in the slow disease subgroup, and a reduction in the number of triple junctions (*P*=0.0072) in intermediate subgroup compared to HC MDMi. No differences in branch junctions were observed in the rapid disease subgroup compared to HC MDMi. The overall evidence demonstrates a clear morphological difference in ALS MDMi, which may be associated with disease progression.

### ALS MDMi reveal abnormal TDP-43 localisation, increased DNA damage and inflammasome formation

A cellular hallmark of ALS is the accumulation of abnormal cytoplasmic TDP-43, where nuclear TDP-43 depletes into the cytosol, forming toxic cytosolic aggregates and inclusions (Neumann et al., 2007; Svahn et al., 2018). Although there is limited understanding of abnormal cytoplasmic TDP-43 localisation in microglia, it has been shown that aberrant TDP-43 regulation can trigger disease specific changes in microglial profiles, such as microglial clearance of amyloid-beta (Aβ), and activated microglial morphology (Paolicelli et al., 2017), where genes associated with phagocytosis are upregulated (Spiller et al., 2018). Further, extracellular TDP-43 proteins promote the toxic production of microglial pro-inflammatory cytokines through nuclear factor-κB (NF-κB), and NLRP3 inflammasome activation in mouse models of ALS (Deora et al., 2020; Zhao et al., 2015). Given that these studies support a toxic role for TDP-43 in microglia, we examined whether TDP-43 pathology was evident in ALS MDMi.

To examine TDP-43 distribution, immunofluorescence was performed. Interestingly, we observed a variety of intracellular TDP-43-positive inclusions including cytoplasmic dashes, spheres, skein-like structures, dots, and round inclusions by immunofluorescence for total TDP-43 (aa 405-414) or phosphorylated (p)TDP-43 (Ser409/410) in ALS MDMi, indicating diverse forms of TDP-43 accumulation in different ALS patients (Fig. 3a-b). The heterogeneous intracellular inclusions were independent of the type of disease progression (slow, intermediate, or rapid), suggesting that there is a cell intrinsic TDP-43 pathology in ALS MDMi, with a range of different TDP-43-positive structures. In support of this, the TDP-43 structures were morphologically similar to inclusions previously reported in post-mortem brain and spinal cord tissues of ALS patients (Brettschneider et al., 2014; Mackenzie et al., 2007; Smethurst et al., 2016). Moreover, TDP-43 aggregation in our cultures was formed under basal conditions and without additional stressors, indicative of *de novo* TDP-43 pathology. Notably, some ALS MDMi appear to have normal nuclear TDP-43 localisation (slow = 33.3%, intermediate = 50%, rapid = 66.7%), indicating a patient-specific difference within disease subgroups. In contrast, HC MDMi presented consistent normal nuclear TDP-43 localisation, and no abnormal TDP-43 aggregates were detected (Fig. 3a). To further examine the cellular distribution of TDP-43, we quantified the ratio of nuclear to cytosolic TDP-43 staining intensities in MDMi and observed that there was significantly increased TDP-43 cytoplasmic staining in ALS compared to HC MDMi (P=0.0002) (Fig. 3c). These data support the involvement of TDP-43 mislocalisation in microglia as a potential pathological feature of ALS.

**Figure 3.**
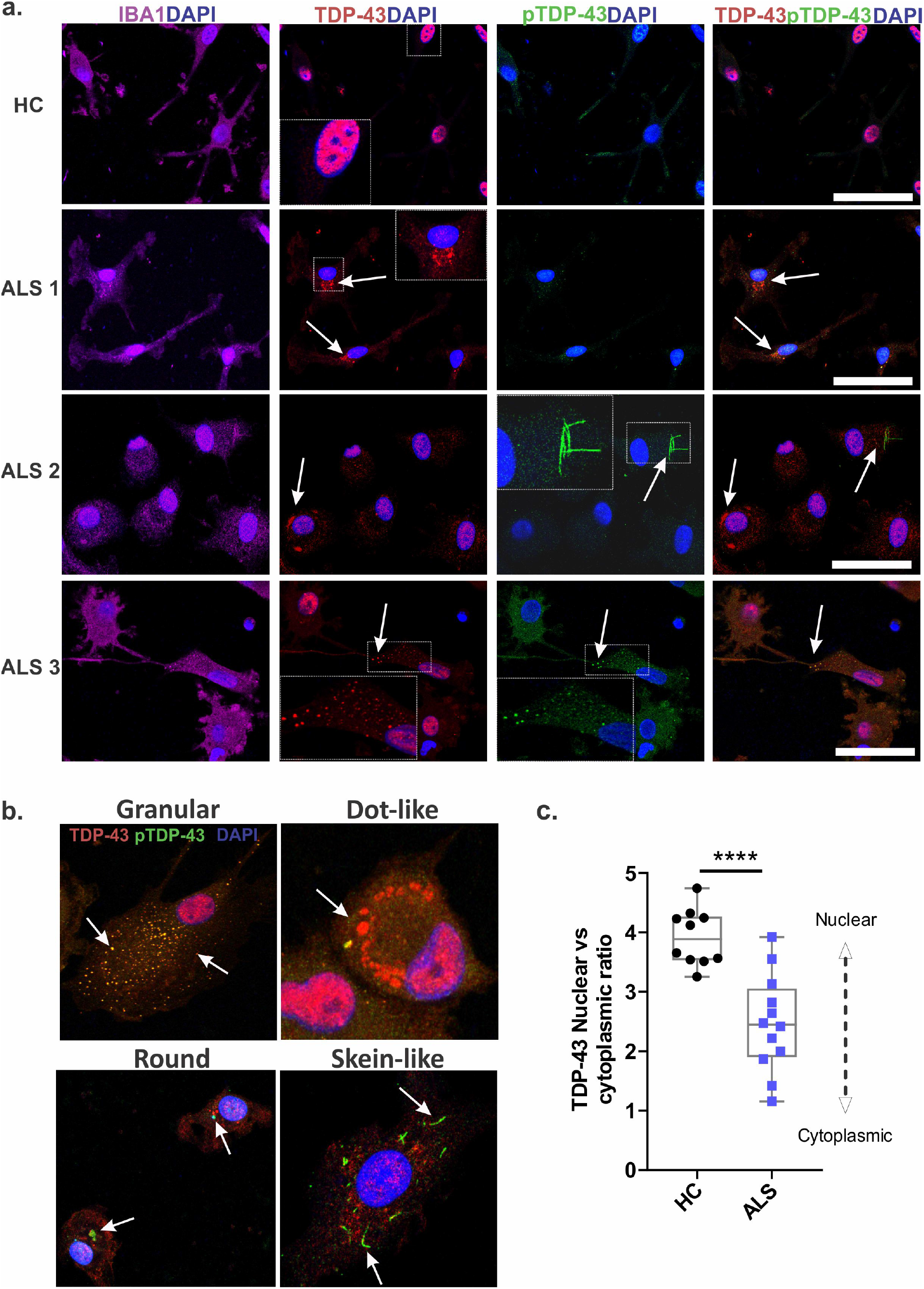
ALS MDMi display abnormal TDP-43 localisation. **(a)** Representative immunofluorescence images of HC and ALS MDMi showing Iba1 (magenta), TDP-43 (red), pTDP-43 (green), and DAPI counterstain (blue). White arrows indicate TDP-43 and/or pTDP-43 inclusions in ALS MDMi. **(b)** White arrows indicate examples of inclusions observed in ALS MDMi including granular, dot-like, round and skein-like. **(c)** Ratio of nuclear to cytoplasmic TDP-43 in HC and ALS MDMi analysed by ImageJ. Significantly increased cytoplasmic TDP-43 observed in ALS compared to HC MDMi. HC: n=10, ALS: n=12. Statistical analysis between two groups was performed using Student’s *t* test. Values are mean ± SEM (***P < 0.001). Scale bars= 50μm.

Given that TDP-43 mislocalisation is involved in the accumulation of DNA damage (Konopka et al., 2020), we evaluated DNA damage in ALS MDMi using an antibody recognising phosphorylated H2AX (γH2AX), which labels DNA double-strand breaks (DSBs). We found significantly increased γH2AX foci positive cells in ALS (30%) compared to HC MDMi (4%) (Fig. S3a,b). Additionally, a subset of ALS MDMi cells (20%) were positive for pan-nuclear γH2AX phosphorylation, while absent in HC MDMi (Fig. S3c). These pan-nuclear γH2AX are formed by clusters of γH2AX foci and are mechanistically and morphologically distinct from γH2AX foci (Meyer et al., 2013). Overall, these results are indicative of increased DNA damage within ALS cells.

Extracellular TDP-43 protein has been shown to activate the NLRP3 inflammasome in microglia from mouse models of ALS (Deora et al., 2020; Zhao et al., 2015). The assembly of the inflammasome complex involves the upregulation of NLRP3 protein, the recruitment of ASC adapter protein, and caspase-1, leading to the cleavage and release of mature IL-1β and IL-18 cytokines and induction of pyroptotic cell death (Schroder & Tschopp, 2010). The NLRP3/ASC/caspase-1 inflammasome dependent activation of the innate immune system has been demonstrated in brain tissues of ALS patients (Johann et al., 2015; Kadhim, Deltenre, Martin, & Sébire, 2016). Given that we observed TDP-43 aggregation/accumulation in ALS MDMi, we next examined for the occurrence of inflammasome activation.

When the inflammasome is activated, cytosolic ASC assembles into a single perinuclear speck structure that co-localises to NLRP3 (Fernandes-Alnemri et al., 2007). Here, we observed the co-localisation of ASC with NLRP3 antibody in a small subset of cells (3%), suggesting an inflammasome activation in ALS MDMi (Fig. S3d). Interestingly, not all ASC foci were colocalised to NLRP3, suggesting that other inflammasomes may be involved. Inflammasome formation is likely to induce pyroptosis in microglia (Deora et al., 2020). Hence, low inflammasome formation observed may reflect the ability to capture inflammasome formation before pyroptosis. No co-localisation of ASC protein and NLRP3 inflammasome was observed in HC MDMi. The activation of NLRP3 inflammasome was also found to be cell-specific and was independent of sex and the disease subgroup (slow, intermediate, and rapid) indicating a dysfunction broadly associated with ALS MDMi. These observations were found under basal conditions and without stressors. Additional studies are needed to examine the full activation of inflammasome formation, including caspase-1 recruitment, processing of IL-1β, and pyroptotic cell death to better understand the role of inflammasome in microglia.

Taken together, these results demonstrate the ability of MDMi to reflect a range of potential pathogenic pathways, which may be the key to elucidating the role of TDP-43 in ALS.

### ALS MDMi reveal altered cytokine expression profiles compared to HCs

We next examined the inflammatory cytokine responses in ALS and HC MDMi. qPCR was used to measure the mRNA expression of classical pro-inflammatory cytokines (*IL-6, IL-8, TNF-α, IL-1β, IL-18)* and alternatively activated cytokines (*IL-10, TGFβ).* Interestingly, we found an upregulation of *IL-8* expression in all ALS MDMi compared to HC MDMi (P=0.0022) (Fig.4, S4). This significant upregulation was observed across each of the disease progression subgroups (slow, P=0.04, intermediate, P=0.0131, rapid, P=0.0034). IL-8 cytokine is known to control migration and activation of microglia and its increase may contribute to, and enhance the inflammatory cascade during disease progression. The observations above corroborate with studies showing elevated levels of IL-8 in cerebrospinal fluid (CSF) and blood of ALS patients compared to healthy individuals (Ehrhart et al., 2015; Kuhle et al., 2009).

**Figure 4.**
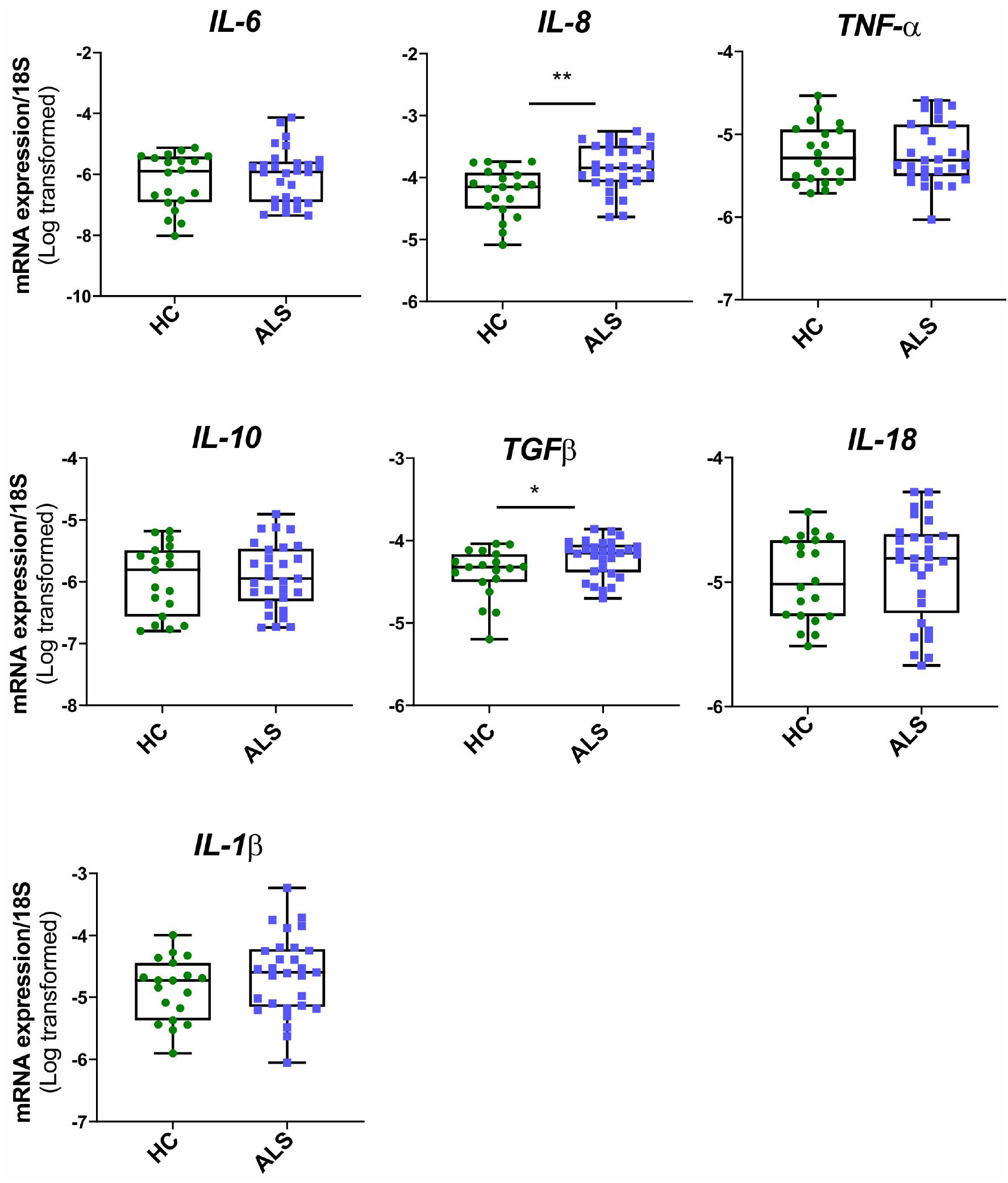
Altered cytokine expression in ALS compared to HC MDMi. Log-transformed mRNA expression of pro-inflammatory cytokines (IL-6, IL-8, TNFα, IL-1β, IL-18) and alternatively activated cytokines (IL-10, TGFβ) in HC and ALS MDMi. HC: n=19, ALS: n=29. Statistical analysis between two groups was performed using Student’s *t* test. Values are mean ± SEM (*P < 0.05, ** *P* < 0.01).

In addition, we found an overall upregulation of *TGFβ* expression in ALS compared to HC MDMi (P=0.0191), where significant upregulation was found in intermediate (P=0.0163) and rapid (P= 0.0192) disease subgroups compared to HC MDMi (Fig 4, S4). Interestingly, no significant difference was observed for *TGFβ* expression in the slow disease subgroup compared to HC (P=0.4311), although the *TGFβ* expression level in the slow disease subgroup was significantly lower compared to intermediate (P=0.0191) and rapid (P=0.0411) subgroups (Fig. S4a). Increased levels of *TGFβ* mRNA expression have been reported to occur in the symptomatic stages of ALS associated with microglial activation and reactive astrogliosis, leading to motor neuron cell loss (Meroni et al., 2019).

There were no obvious changes in the expression of other cytokines (*IL-6, TNF-α, IL-10, IL-1β, IL-18*). Heterogeneity was present in cytokine expression across ALS and HC individuals, suggesting that further investigation of cytokine expression at an individual level may allow for stratification of disease-associated changes.

### Sex-specific variability of cytokine expression in ALS and HC MDMi

Given that ALS affects males more than females (McCombe & Henderson, 2010), we compared ALS and HC MDMi according to sex. Males represented 45% (HC), 66% (slow), 40% (intermediate) and 55.6% (rapid) of subjects in each group. Interestingly, we found significantly increased *TNFα* expression in males (m) compared to females (f) in the slow and rapid disease subgroups (Fig. 5). Elevated levels of *TNFα* has been found in the serum and in the CSF of ALS patients, which further correlates with disease progression and severity (Fukazawa, Tsukie, Higashida, Fujikura, & Ono, 2013; Poloni et al., 2000). However, little is known regarding the sex-specific alteration of *TNFα* expression in ALS, which should be explored in future studies (Cereda et al., 2008; Hu et al., 2017).

**Figure 5.**
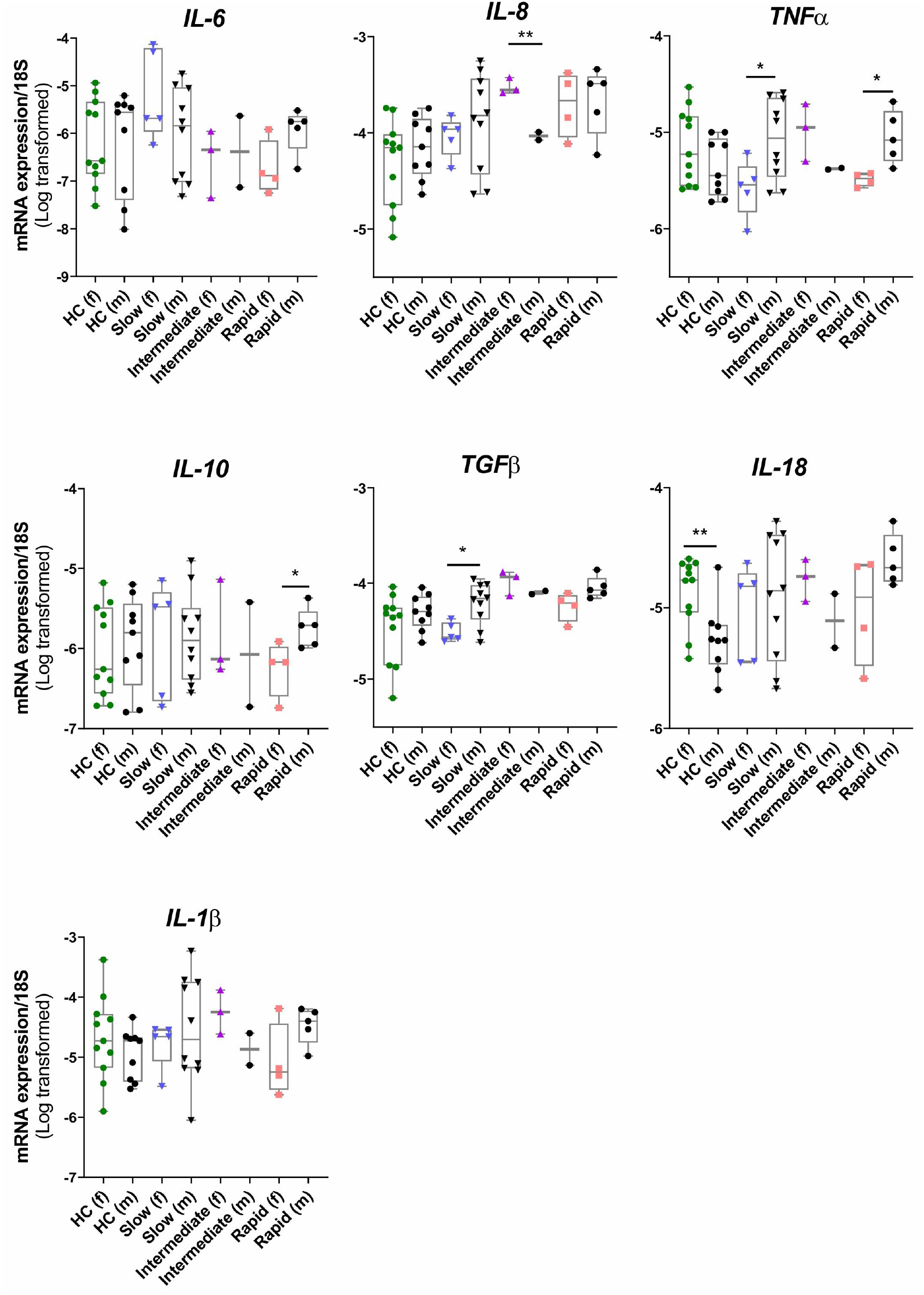
Sex-specific cytokine expression profiles in ALS subgroups and HC MDMi. Log-transformed mRNA expression of pro-inflammatory cytokines (IL-6, IL-8, TNFα, IL-1β, IL-18) and anti-inflammatory cytokines (IL-10, TGFβ) in HC and ALS subgroups MDMi. HC (f): n=11, HC (m): n=9, slow (f): n=5, slow (m): n=10, intermediate (f): n=3, intermediate (m): n=2, rapid (f): n=4, rapid (m): n=5. Statistical analysis between two groups was performed using Student’s *t* test and between multiple groups using one-way ANOVA. Values are mean ± SEM (*P < 0.05, ** *P* < 0.01). f: females, m: males.

The change in expression of *TGFβ* mRNA observed in ALS cases (Fig. 4) correlated with increased expression in males compared to females, reaching significance in the slow disease subgroup (Fig. 5). Further, significantly increased *IL-10* mRNA expression was observed in males in the rapid disease subgroups, while *IL-1β* and *IL-18* expression was elevated but not significantly altered compared to female patients. *IL-18* was also found to be significantly lower in HC males compared to HC females. Conversely, an upward trend of *IL-18* expression was observed in males in slow and rapid disease subgroups.

### ALS MDMi have impaired phagocytosis

Phagocytosis is an essential function of microglia, and is reportedly altered in ALS (Haukedal & Freude, 2019). We therefore examined the phagocytic ability of ALS MDMi by adding a pH-sensitive dye phagocytic probe (pHrodo-labelled *E.coli* particles) into MDMi cultures and imaged each hour using the IncuCyte ZOOM live imaging platform. pHrodo dyes are non-fluorescent in media and phagocytic function will present as an increase in pHrodo-specific fluorescence under conditions of low pH such as incorporation into maturing phagosomes and endocytic compartments.

We observed significantly decreased phagocytic uptake of labelled *E.coli* particles in ALS compared to HC MDMi, as determined by measurement of total pHrodo-labelled particle area per cell (2.6-fold reduction) (Fig. 6a-c, S5a-d), which was further confirmed by decreased area of labelled *E.coli* particle uptake per cell (3.4-fold reduction) (Fig. 6d). Moreover, phagocytosis impairment revealed a trend associated with increasing disease severity with a 60% reduction in phagocytic activity in the slow disease subgroup (P=0.015), 74% reduction in the intermediate disease subgroup (P=0.0012), and 79% reduction in the rapid disease subgroup (P>0.0001) compared to HC MDMi (Fig. S6f, representative pictures S5e).

**Figure 6.**
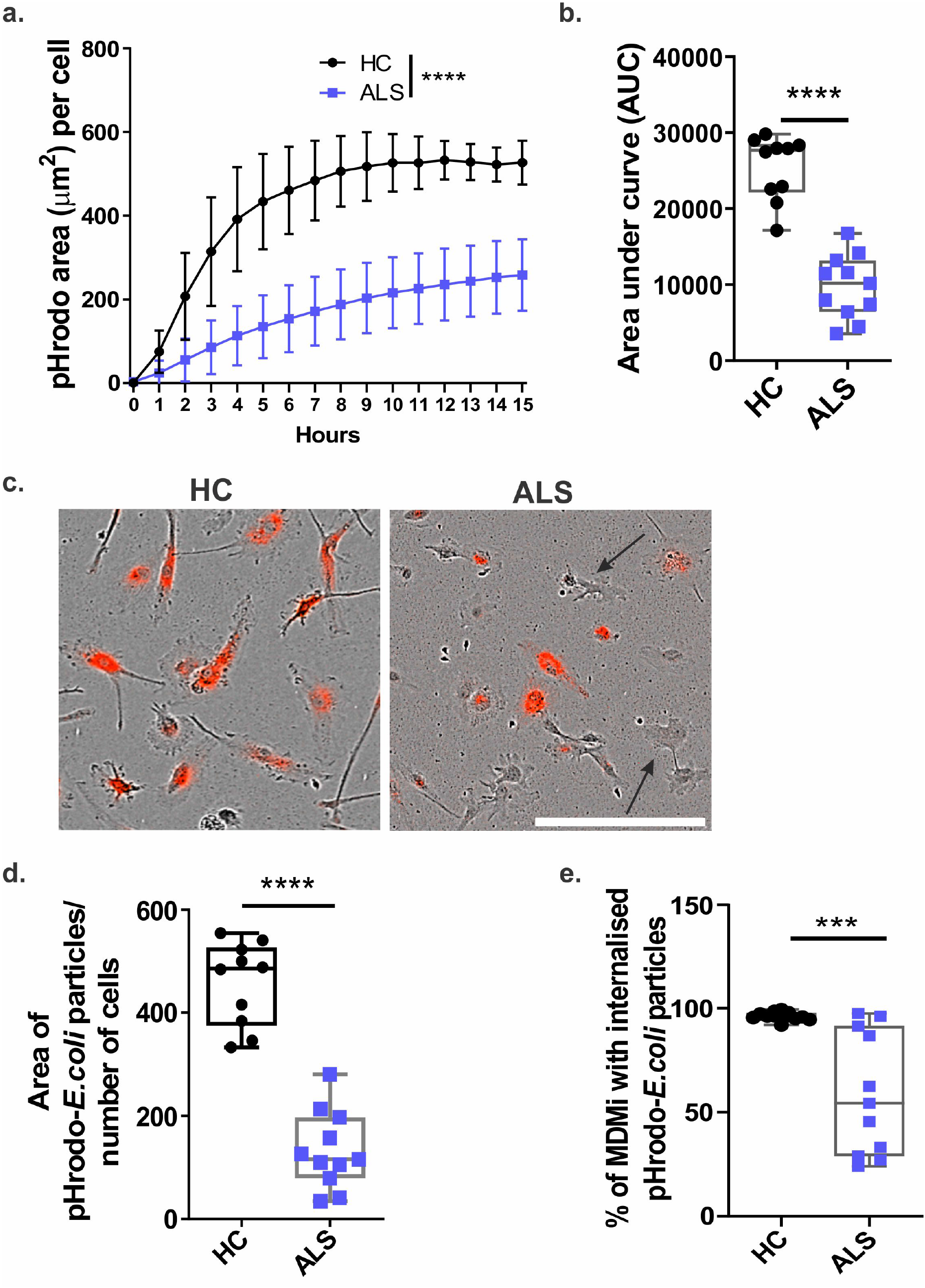
Impaired phagocytosis in ALS compared to HC MDMi. **(a)** Quantification of phagocytosis by pHrodo-labelled *E.coli* particles over 15 hrs using live imaging in HC and ALS. **(b)** Uptake of pHrodo-labelled *E.coli* particles in HC and ALS MDMi was quantified using area under the curve. **(c)** Representative images of HC and ALS MDMi uptake of pHrodo-labelled *E.coli* particles (red). Black arrows indicate impaired phagocytosis in specific cells in ALS MDMi. **(d)** Area of pHrodo-labelled *E.coli* particles normalised over cell number, quantified using IncuCyte ZOOM in-built software. **(e)**. Percentage of cells that phagocytose particles in HC and ALS MDMi. HC: n=10, ALS: n=11. n=200 cells per patient or individual. Statistical analysis between two groups was performed using Student’s *t* test and between multiple groups using one-way ANOVA. Values are mean ± SEM (***P < 0.001, **** *P* < 0.0001). Scale bar = 50μm.

**Figure 7.**
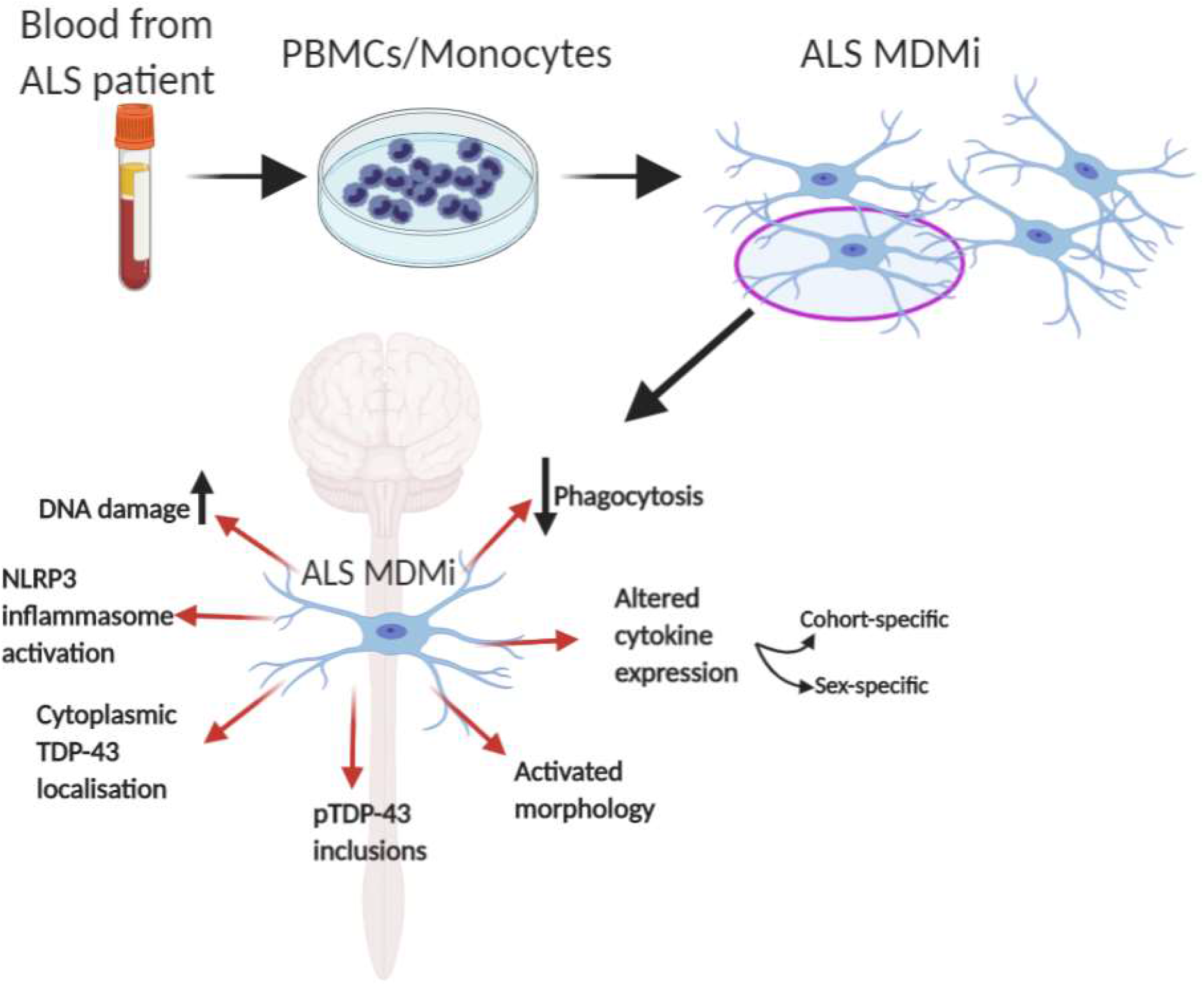
Characterisation of ALS MDMi and pathological pathways for potential microglia targeted therapy. Blood monocytes derived from ALS patient PBMCs were successfully cultured to microglia (monocyte-derived microglia, MDMi). ALS MDMi recapitulated hallmarks of ALS pathology, including cytoplasmic TDP-43 localisation and phosphorylated (p)-TDP-43 inclusions. A range of abnormalities including microglial activation, altered cytokine expression, and decreased phagocytosis was observed in ALS MDMi compared to HC MDMi. MDMi model is highly suited to investigate patient heterogeneity, screening of patients for clinical trials, and providing a basis for automated drug screening platforms in ALS and other neurodegenerative diseases.

This impairment in phagocytic function appeared to be cell-specific, with some individual MDMi showing a strong impairment while other cells in the same culture appeared relatively normal (Fig. S5f-g, representative pictures S5e). We examined this further and observed that 59% of ALS MDMi cells had internalised labelled-E.coli particles compared to 96% of HC MDMi cells (P=0.0003, Fig. 6e, representative picture 6c). Within the ALS disease subgroups, phagocytosis again showed greater impairment in MDMi from the rapid disease subgroup (70% reduction in number of cells showing uptake, P>0.0001) followed by intermediate (40% reduction, P=0.0012) and slow (12% reduction, P=0.015) disease subgroups compared to HC MDMi.

Taken together, these findings indicate that a sub-population of ALS MDMi have dysfunctional phagocytic function and this correlates with the rate of disease progression.

## Discussion

Neuroinflammation is a hallmark of ALS, which is mediated by activated glial cells and infiltrating immune cells from the periphery leading to the subsequent neuronal cell death in the brain and spinal cord (Brettschneider et al., 2012; Endo et al., 2015; Turner et al., 2004). The complex role of the peripheral and the CNS immune dysregulation in the pathogenesis of ALS is not well understood due to the lack of a cellular model that is able to recapitulate the clinical heterogeneity in patients over the course of disease progression. Evidence for the role of activated microglia in ALS has stemmed largely from human post-mortem analysis, and animal models that represent rare familial forms of the disease. Due to the difficulty of studying the CNS in living people with ALS, few studies have investigated inflammatory events at earlier time points in sporadic cases of disease. Here, we have presented for the first time, a patient-derived *in vitro* model (MDMi) to study microglia-like cells from living ALS patients to increase our understanding of the functional phenotype of microglia and their potential role in ALS pathogenesis. More importantly, the MDMi model remains in its myeloid lineage throughout differentiation, retaining specific epigenetic or splicing patterns in myeloid cells that are patient-specific and are reflective of disease changes.

MDMi most resemble primary fetal brain microglia, displaying enriched microglial genes (*P2RY12, C1QA, MERTK, PROS1, GPR34* and *GAS6)* (O. Butovsky et al., 2012; Sellgren et al., 2017). Less enriched microglial markers were observed in isolated adult *ex vivo* microglia (serum-starved for 24hrs), in cultured brain *ex vivo* primary microglia (Gosselin et al., 2017), and in an immortalised microglial cell line (Sellgren et al., 2017). One explanation for the differential gene expression between resident microglia and *ex vivo* or cultured microglia could be the lack of cell-cell interaction with neurons and other glial cells that drives gene expression. This has been supported by studies showing rapid changes to microglia transcriptomes after isolation from mouse brain (Gosselin et al., 2017). Thus, future research into microglia-related ALS pathology would benefit from co-culturing MDMi with other cell types found within the CNS in order to better recapitulate a brain microenvironment.

A substantial reduction in the phagocytic capacity was observed in ALS MDMi, which was further shown to occur in a heterogeneous manner. A proportion of MDMi generated from each patient revealed low levels of phagocytic activity while other MDMi from the same patient had normal phagocytosis (Fig. 6). This heterogeneous dysfunction was seen across all of the ALS subgroups (slow, intermediate, and rapid disease progression), but was exacerbated in rapid compared to intermediate and slow disease progression subgroups, suggestive of an association between microglia phagocytic impairment and rate of disease progression. The molecular basis of impaired phagocytic capacity in patient-specific MDMi within the same culture is unclear, but could be related to the monocyte sub-population (classical: CD14^++^/CD16^-^, intermediate: CD14^+^/CD16^+^, and non-classical: CD14^+^/CD16^++^) and its inflammatory state over the course of disease progression (O. Butovsky et al., 2012; Du et al., 2020; McGill et al., 2020; Zhao et al., 2017; Zondler et al., 2016). Hence, it would be of interest to examine if the different monocyte sub-populations are reflected in MDMi by enrichment of monocyte sub-populations prior to differentiation, and/or performing single cell RNA-sequencing analysis of ALS MDMi cultures in future studies.

TDP-43 proteinopathies are predominantly characterised by degenerating neurons, astrocytes and muscle fibres, which are subsequently phagocytosed by microglia (Spiller et al., 2018; Svahn et al., 2018). Interestingly, our study revealed that ALS MDMi recapitulated *de novo* TDP-43 aggregates and increased phospho-TDP-43 immunoreactivity, which are major hallmarks of ALS. These observations further support the growing evidence for TDP-43 pathology in microglia (Paolicelli et al., 2017), and other non-neuronal cell types including skin fibroblasts (Riancho et al., 2020), circulating lymphomonocytes, and in monocytes/macrophages from a subgroup of ALS patients (De Marco et al., 2017; De Marco et al., 2011). Our study is the first to report the appearance of abnormal TDP-43 pathology in microglia generated from people living with ALS.

Paolicelli et al. demonstrated a role for TDP-43 in regulating microglial phagocytosis, where increased levels of CD68, a microglial activation marker, was observed in ALS patients with TDP-43 pathology compared to ALS patients with no TDP-43 pathology. Further, knockdown of TDP-43 protein expression in a BV2 mouse microglial cell line, enhanced phagocytic activity. These findings support a role for TDP-43 in regulation of microglial phagocytosis. Interestingly, we did not observe a strong relationship between impaired phagocytosis in ALS MDMi and TDP-43 mislocalisation, suggesting that other key factors in ALS microglia may be associated with these pathological changes in microglia function, which needs to be further elucidated. Nevertheless, abnormal TDP-43 pathology in ALS microglia may impair their phagocytic ability, alter TDP-43 expression in neuronal and glial cells, perpetuating a selfdriving neuroinflammatory loop. Overall, our findings are consistent with a role for dysfunctional microglia phagocytosis in TDP-43 proteinopathies.

Microglia activation can result in a neurotoxic or neuroprotective phenotype depending on the degree and type of inflammation in ALS (Beers & Appel, 2019). We show here that the addition of exogenous TDP-43 recombinant protein in healthy MDMi was able to significantly upregulate classical pro-inflammatory cytokine expression and downregulate antiinflammatory or alternative activation cytokines (the latter, are believed to be associated with a more protective outcome for neurons). In keeping with this, MDMi from ALS showed an upregulation of *IL-8* (pro-inflammatory), and *TGFβ* (potentially protective), while no significant changes in the expression levels of *IL-6, IL-10, IL-1β, IL-18* were observed under the same conditions. A possible explanation for the lack of substantial changes across most cytokine expression in the ALS cohort could be due to patient heterogeneity and limited sample size. Nevertheless, increased levels of IL-8 protein in serum and CSF has been observed in ALS patients (Ehrhart et al., 2015; Mitchell et al., 2009).

However, it is unclear if cytokine responses are directly associated with TDP-43 pathology. RNA-sequencing of nuclear and cytoplasmic brain fractions from transgenic mice with abnormal nuclear TDP-43 localisation demonstrated altered inflammatory pathways (Amlie-Wolf et al., 2015), indicating that abnormal sub-cellular localisation of TDP-43 may affect inflammatory cytokine generation and/or secretion. However, we did not observe a direct correlation between abnormal TDP-43 localisation and cytokine expression, suggesting that the relationship between these changes is complex and may require further evaluation in larger cohorts.

As a consequence of inflammation within the CNS, the morphology of microglial cells changes from a ramified to amoeboid morphology (activated state), with enlarged soma and decreased branch length (Frank-Cannon et al., 2009). We have shown here that cell processes were shorter in ALS compared to HC MDMi as evident by reduced branch length and end-points, indicative of microglial activation. This was observed in slow, intermediate and rapid ALS disease subgroups although there were no disease subgroup-specific morphological changes. These results are consistent with studies that have examined ALS spinal cord tissue and demonstrated enlarged cell processes in amoeboid-like CD68-positive microglia associated with extensive neuronal loss (Brettschneider et al., 2012). There was no correlation between abnormal MDMi morphology and TDP-43 pathology, age, sex, or disease progression indicating that morphological changes in ALS MDMi are a consistent factor across the disease and patient spectrum.

We further stratified the ALS cohort into disease subgroups and sex, and observed the upregulation of TNFα in males compared to females in slow, and rapid disease progression subgroups. TNFα expression is regulated by two main signalling receptors, p55 TNFα receptor 1 (TNFR1), and p75 TNFα receptor 2 (TNFR2), which either promote a neurotoxic or neuroprotective role respectively (Tortarolo et al., 2017). Clinical data demonstrate that both receptors were found to be upregulated in spinal cord tissues of ALS patients (Tortarolo et al., 2017), indicating a dysregulation of these receptors in disease pathology. The contribution of sex to TNFR1 and TNFR2 regulation in ALS has yet to be reported, but has been demonstrated in other studies (Miller, Bonn, Franklin, Ericsson, & McKarns, 2015; Wang, Crisostomo, Markel, Wang, & Meldrum, 2008). Moreover, differential expression of TNFR1 and TNFR2 have been found in monocyte sub-populations (Hijdra, Vorselaars, Grutters, Claessen, & Rijkers, 2012), supporting the diverse role of monocyte sub-populations that may contribute to ALS MDMi pathology.

We show that MDMi from ALS patients displayed increased γH2AX foci and pan-nuclear γH2AX phosphorylation in ALS MDMi compared to HC MDMi under basal conditions. This result is in line with other reports showing that ALS tissues exhibit increased DNA damage foci, likely mediated by oxidative and replication stress, as recently substantiated in studies of different ALS cell models (Konopka et al., 2020; Mitra et al., 2019; Riancho et al., 2020). Although it has been shown that TDP-43 is recruited less efficiently to DNA damage foci leading to unresolved DDR (Mitra et al., 2019), further studies are needed to examine DDR signalling pathways in ALS MDMi. However, it remains a possibility that unresolved DDR can lead to the accumulation of single-stranded DNA (ssDNA) and dsDNA, which has been observed in the cytoplasm and nucleus of motor neurons, leading to subsequent cell death or inflammation (Martin et al., 2007; Quek et al., 2017). The activation of the innate immune response triggered by NLRP3, is by far the most studied inflammasome implicated in neurodegenerative disease (Heneka, Kummer, & Latz, 2014), yet the role of NLRP3 inflammasome in ALS remains elusive. Given that NLRP3 and ASC were expressed together in ALS MDMi suggest inflammasome activation, however this finding needs to be confirmed by the presence of active caspase 1 and the release of mature IL-1β and IL-18 for complete inflammasome activation. Additionally, ASC positive but NLRP3 negative ALS MDMi cells were observed, indicating that other inflammasome may be involved in microglia mediated inflammatory response. Nevertheless, elevated levels of caspase 1, IL-1b and IL-18 cytokine have been reported in serum and spinal cord tissues of ALS, supporting the role in inflammasome in ALS pathology (Johann et al., 2015). Taken together, the interplay between TDP-43 mislocalisation, neuroinflammation and DNA-damage is a complex process. For instance, (1) dysregulation of TDP-43 can lead to impaired DNA damage (Mitra et al., 2019), DDR, (Konopka et al., 2020), and (2) inflammation (Zhu, Cynader, & Jia, 2015). (3) Unresolved DNA damage and aberrant cytosolic DNA leading to impaired DDR signalling contributes to neuroinflammation (Quek et al., 2017). (4) Persistent neuroinflammation results in oxidative stress that can, in turn, promote cytosolic TDP-43 aggregation and exacerbate DNA damage (Correia, Patel, Dutta, & Julien, 2015).

There is currently no effective treatment for the progressive neurodegeneration in ALS, hence new treatment options are urgently needed. However, a limiting factor in elucidating the pathological role of microglia over the course of ALS disease progression is the difficulty in obtaining relevant tissue samples from living ALS patients. In this regard, we have shown here that MDMi generated from patient blood monocytes provides a readily accessible model that displays key microglial features including ramified morphology, phagocytosis, and inflammatory responses. Further, we show that the MDMi model is able to reflect patient heterogeneity, which potentially enables personalised therapeutics and a clinically relevant drug screening platform for microglial targeting compounds. We show that ALS MDMi recapitulate key aspects of ALS neuropathology including TDP-43 aggregation, activated microglial morphology, DNA damage, cytokine dysregulation, and impaired phagocytosis. We also identified important cell- and disease subgroup-specific differences in phagocytic function that warrant further examination for cell-specific changes occurring at the molecular level. Further studies using this robust, cost-effective and reproducible approach of generating patient microglia could help fast-track new personalised therapeutic treatments for ALS patients.

## Author contributions

H.Q and A.R.W conceived and designed the experiments. T.C, A.N, V.L.B coordinated blood collection from the ALS clinical Research Centre in Palermo, Italy. Y.S, C.C.G, M.K.L coordinated blood collection from the PISA study. H.Q and C.C.L generated experimental data and figures. H.Q, C.C.L, R.S, T.H.N, L.E.O, T.L.R, Y.C.L, and A.R.W analysed experimental data. H.Q and A.R.W wrote the manuscript, with input from all authors.

## Acknowledgement

The authors would like to thank the volunteers who have donated blood to this project from QIMRB, and participants recruited through the ALS clinical at University of Palermo and the PISA study. We also thank the QIMR Berghofer Medical Research Institute Microscopy, Sample Processing team and the QIMR statistical team for their assistance. We thank Dr Wayne A. Schroder for his advice on dendritic cells and acknowledge the coordinators of the PISA study, Professor Michael Breakspear, Jessica Adsett and Natalie Garden.

## Supplementary Materials

**Supplementary Table 1: List of qPCR primers**

**Supplementary Figure 1. Gene expression analysis of MDMi with isolated monocytes, monocyte-derived macrophages (MDMa), and monocyte-derived dendritic cells (MDDCs).**

**Supplementary Figure 2. Morphology of ALS MDMi.**

**Supplementary Figure 3. ALS MDMi display increased DNA damage, and NLRP3 inflammasome formation.**

**Supplementary Figure 4. Altered cytokine expression in ALS disease subgroups compared to HC MDMi.**

**Supplementary Figure 5. Impaired phagocytosis in ALS MDMi.**

---

**Supplementary Table 1.**
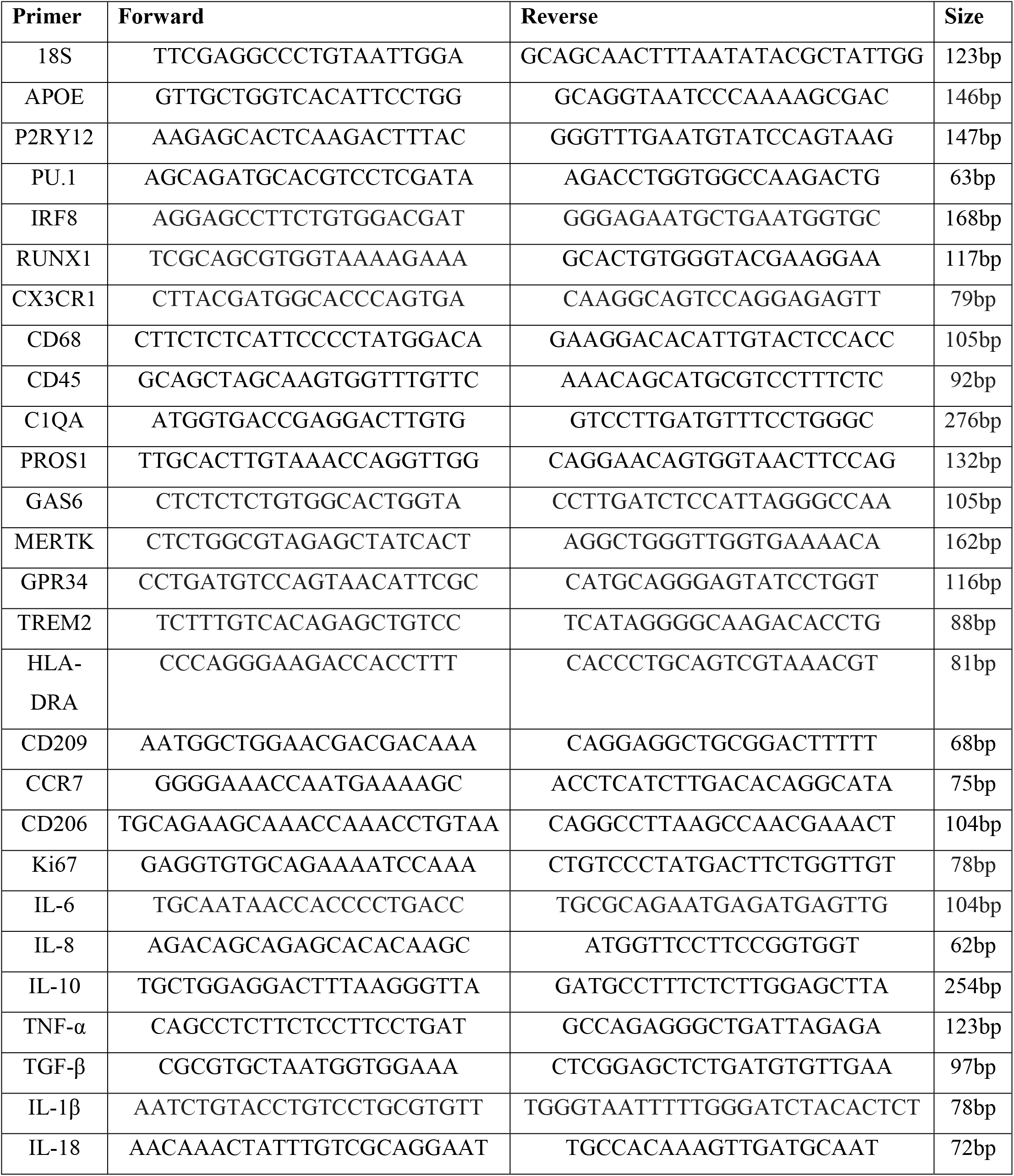
List of qPCR primers

**Supplementary Figure 1.**
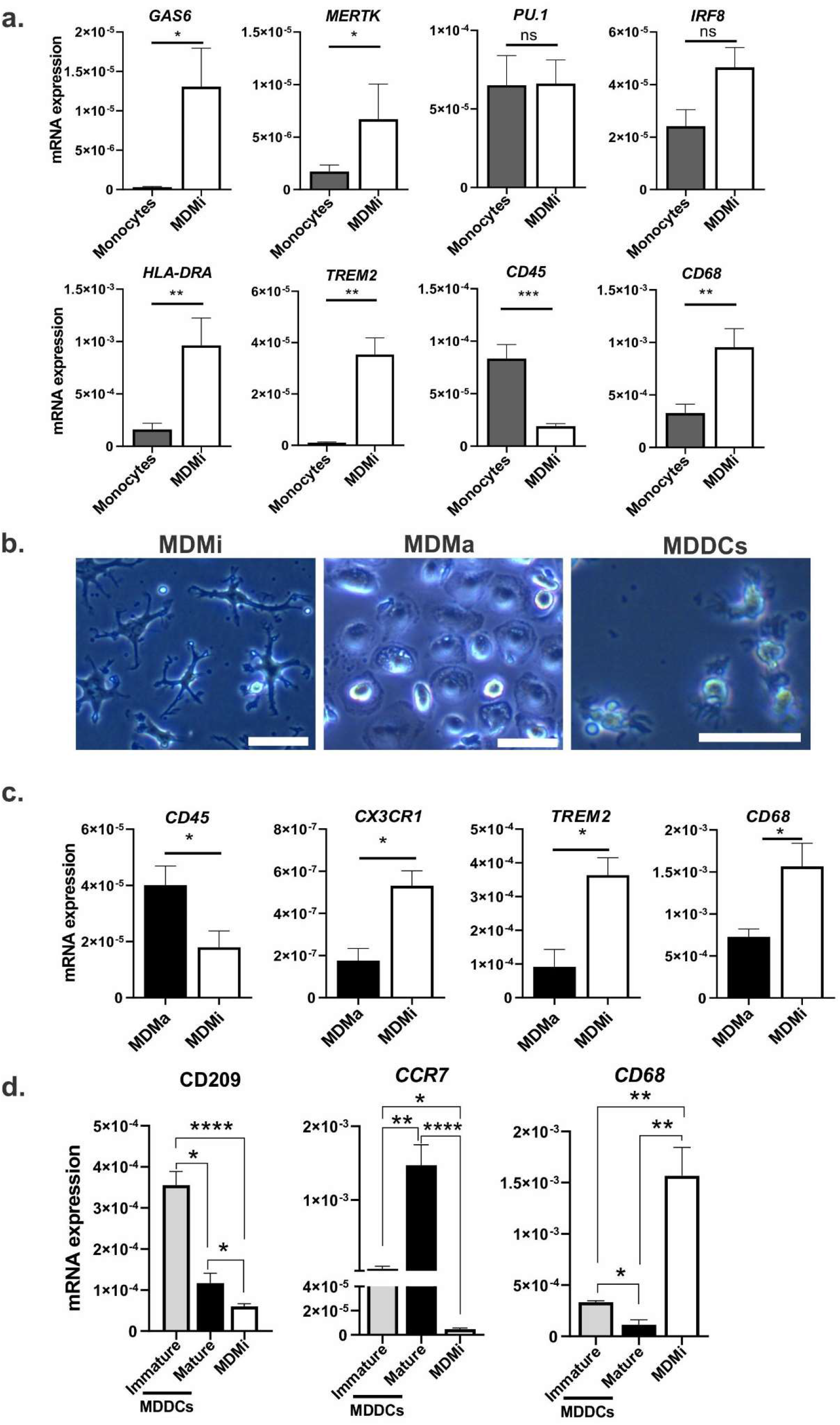
Gene expression analysis of MDMi with isolated monocytes, monocyte-derived macrophages (MDMa), and monocyte-derived dendritic cells (MDDCs). MDMi, MDMa, and MDDCs were generated from monocytes isolated from the same healthy young volunteers (<40 years of age) to examine gene changes. **(a)** Gene expression levels of well-known microglial genes such as *GAS6, MERTK, PU.1, IRF8, HLA-DR, TREM2,* and *CD68* between isolated monocytes and MDMi. A monocyte/macrophage gene, *CD45* confirmed a monocyte phenotype. (Monocytes and MDMi from <40 years of age: n=5). **(b)** Representative phase contrast pictures of MDMi (Day 14), MDMa (Day 14), and MDDCs (Day 9). **(c)** Gene expression levels of well characterised microglial genes *TREM2, CX3CR1* and *CD68,* and monocyte/macrophage gene, *CD45* between MDMa and MDMi. (MDMa and MDMi from <40 years of age: n=5). **(d)** Bar graph showing gene expression levels of dendritic cell markers *CD209* and *CCR7* and microglial marker, *CD68* between MDDCs and MDMi. (MDDCs and MDMi from <40 years of age: n=5). Statistical analysis between two groups was performed using Student’s *t* test and between multiple groups using one-way ANOVA. Values are mean ± SEM (*P < 0.05, ** *P* < 0.01, *** *P* < 0.001, **** *P* < 0.0001). Scale bars= 50μm.

**Supplementary Figure 2.**
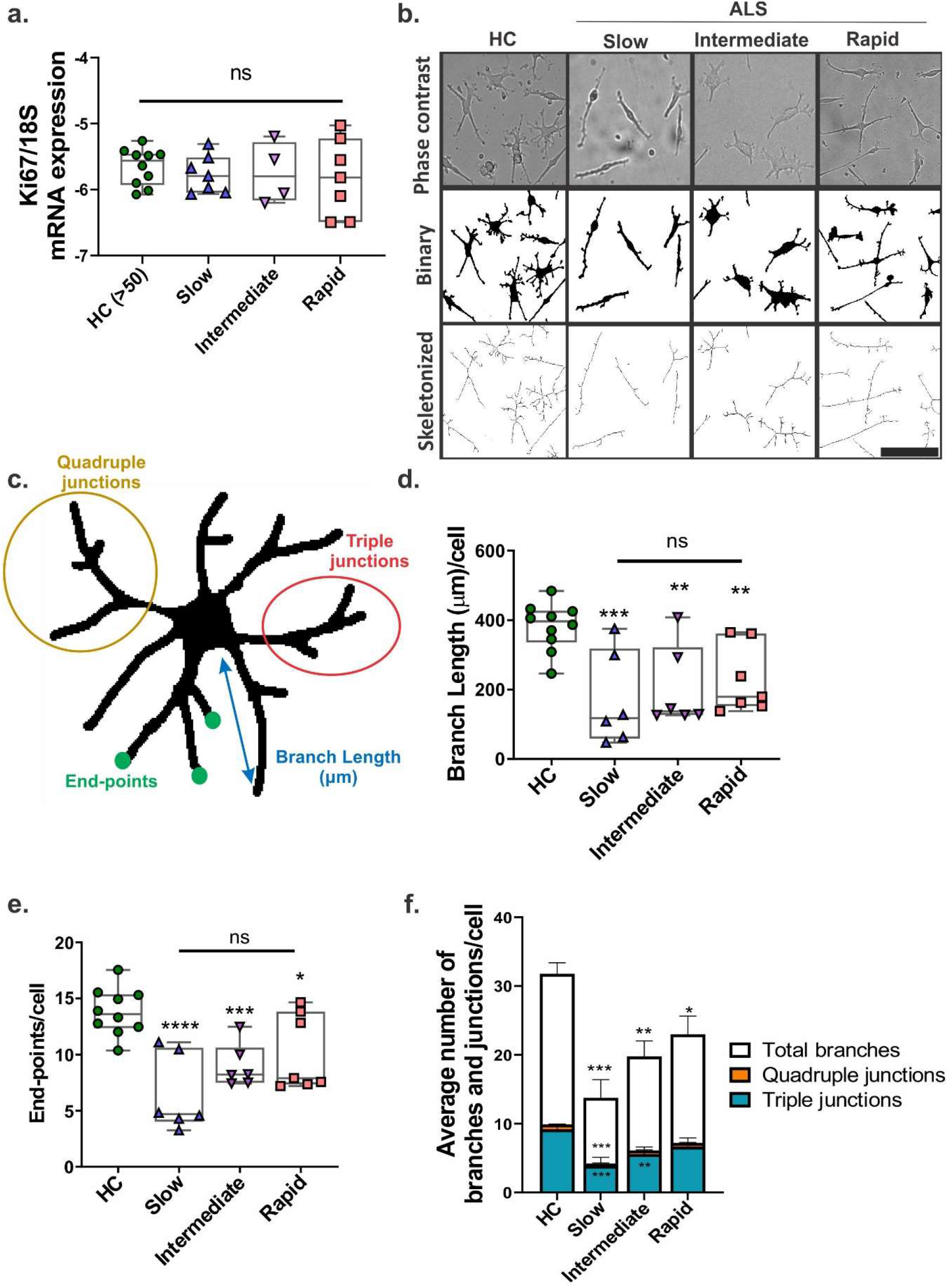
Morphology of ALS MDMi. **(a)** Relative expression of Ki67 mRNA normalised to housekeeping gene, 18S, was examined in HC, slow, intermediate, and rapid ALS MDMi subgroups. No differences were observed between HC and ALS cohort or between ALS MDMi subgroups; HC: n=10, Slow: n=7, Intermediate: n=4, Rapid: n=7. **(b)** Representative images of analysed MDMi morphology in ALS subgroups using ImageJ. Phase contrast images were adjusted to 16-bit and thresholded to create a mask for MDMi cells. Images were then converted to binary and analysed using the AnalyzeSkeleton plugin to obtain various microglial branch information. **(c)** Schematic example of a skeletonised MDMi showing branch length (μm), end-points, triple junctions (junctions with exactly 3 branches), and quadruple junctions (junctions with exactly 4 branches). **(d)** Microglial branch length of HC and ALS MDMi subgroups were analysed by ImageJ and normalised against cell number. Significant reduction of microglial branch length was found in all ALS subgroups compared to HC MDMi, where the slow disease subgroup displayed marked reduction followed by intermediate and rapid compared to HC MDMi. HC: n=10, Slow: n=6, Intermediate: n=6, Rapid: n=7. **(e)** Microglial end-points of HC and ALS MDMi subgroups were analysed by ImageJ and normalised against cell number. Significant reduction of end-points was found in all ALS subgroups compared to HC MDMi. No differences were observed across ALS subgroups. HC: n=10, Slow: n=6, Intermediate: n=6, Rapid: n=7. **(f)** Average number of microglial branches and junctions of HC and ALS MDMi were analysed by ImageJ and normalised against cell number. Significant reduction of total branches was observed in all ALS subgroups compared to HC MDMi. Significant reduction of microglial triple and quadruple junctions was observed in slow disease subgroup compared to HC MDMi, while a reduction of triple junction was observed in intermediate disease subgroup compared to HC MDMi. No difference in microglial branch junctions were observed in rapid disease subgroup. HC: n=10, Slow: n=6, Intermediate: n=6, Rapid: n=7. Statistical analysis between two groups was performed using Student’s *t* test and between multiple groups using one-way ANOVA. Values are mean ± SEM (*P < 0.05, ** *P* < 0.01, *** *P* < 0.001, **** *P* < 0.0001). Scale bar= 50μm.

**Supplementary Figure 3.**
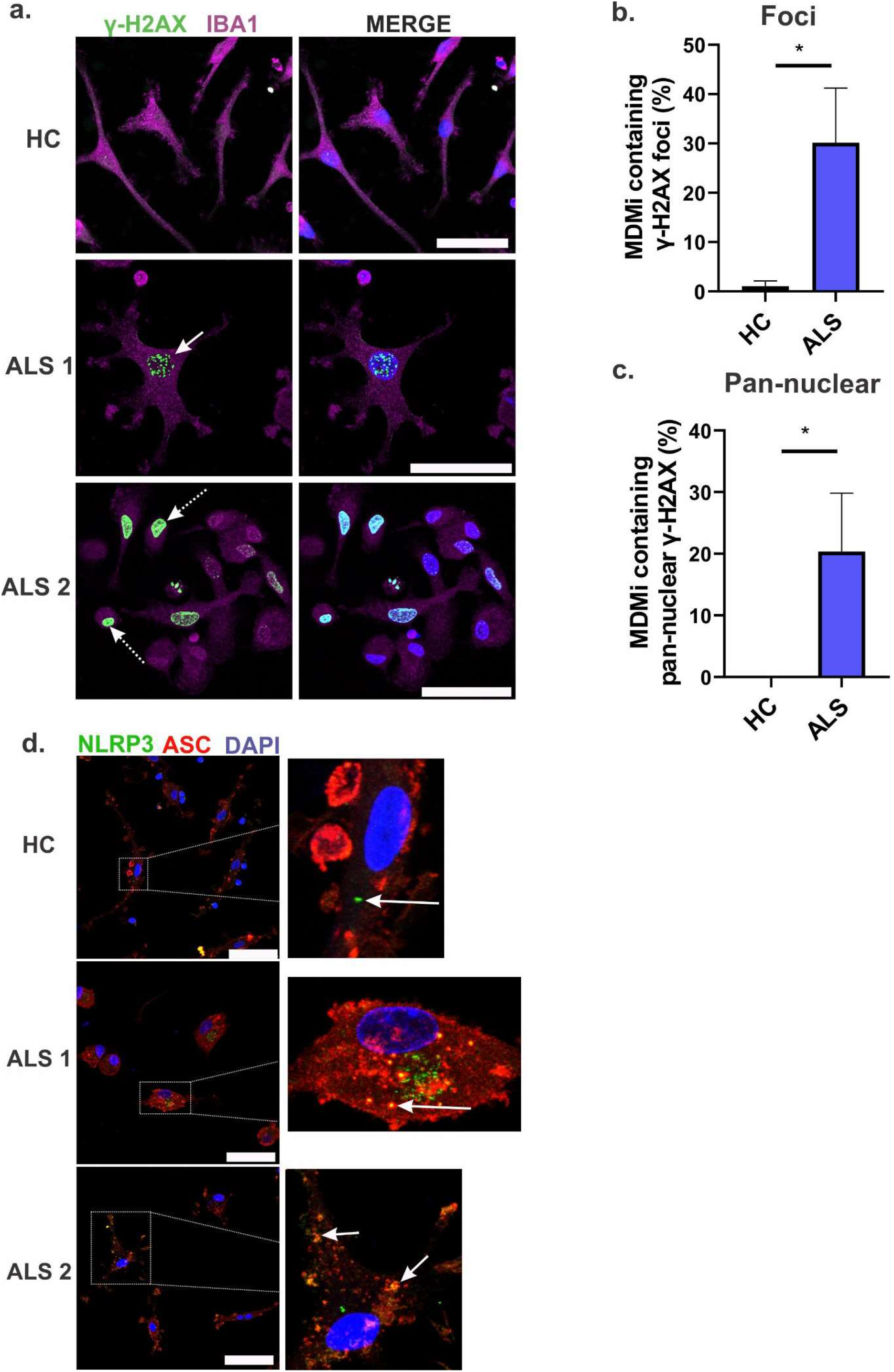
ALS MDMi display increased DNA damage, and NLRP3 inflammasome formation. **(a)** Representative immunofluorescence images of ALS MDMi showing γH2AX (green), Iba1 (magenta), and DAPI counterstain (blue). White solid arrow indicates γH2AX foci, and dotted white arrows indicate pan-nuclear γH2AX staining in ALS MDMi. **(b)** Increased percentage of MDMi containing γH2AX foci in ALS compared to HC MDMi, n=60 cells. **(c)** Increased percentage of MDMi containing pan-nuclear γH2AX staining in ALS compared to HC MDMi, n=60 cells. **(d)** Representative immunofluorescence images of HC and ALS MDMi showing NLRP3 (green), ASC (red) and DAPI counterstain (blue). White solid arrow showing NLRP3 in HC that does not co-localise with ASC, while ASC speck was co-localised with NLRP3 in ALS MDMi suggesting inflammasome formation. HC: n=3, ALS: n=3. Statistical analysis between two groups was performed using Student’s *t* test and between multiple groups using one-way ANOVA. Values are mean ± SEM (*P< 0.05). Scale bars= 50μm. ASC: apoptosis-associated speck-like protein containing a CARD; NLRP3: NLR family pyrin domain containing 3.

**Supplementary Figure 4.**
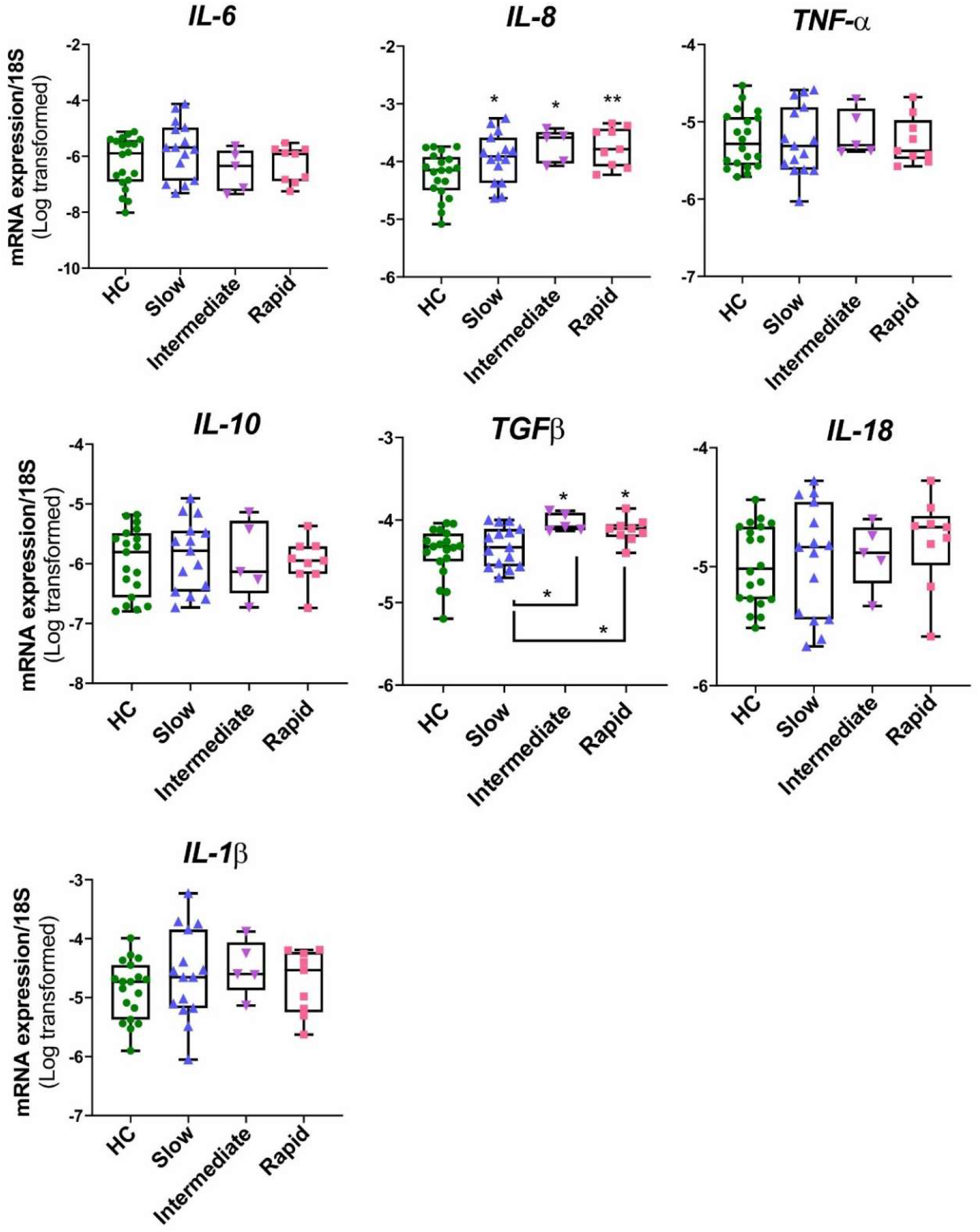
Altered cytokine expression in ALS disease subgroups compared to HC MDMi. Log-transformed mRNA expression of pro-inflammatory cytokines (IL-6, IL-8, TNFα, IL-1β, IL-18) and alternatively activated cytokines (IL-10, TGFβ) in HC and ALS subgroups MDMi. HC: n=19, slow: n=15, intermediate: n=5, rapid: n=9. Statistical analysis between two groups was performed using Student’s *t* test and between multiple groups using one-way ANOVA. Values are mean ± SEM (*P < 0.05, ** *P* < 0.01).

**Supplementary Figure 5.**
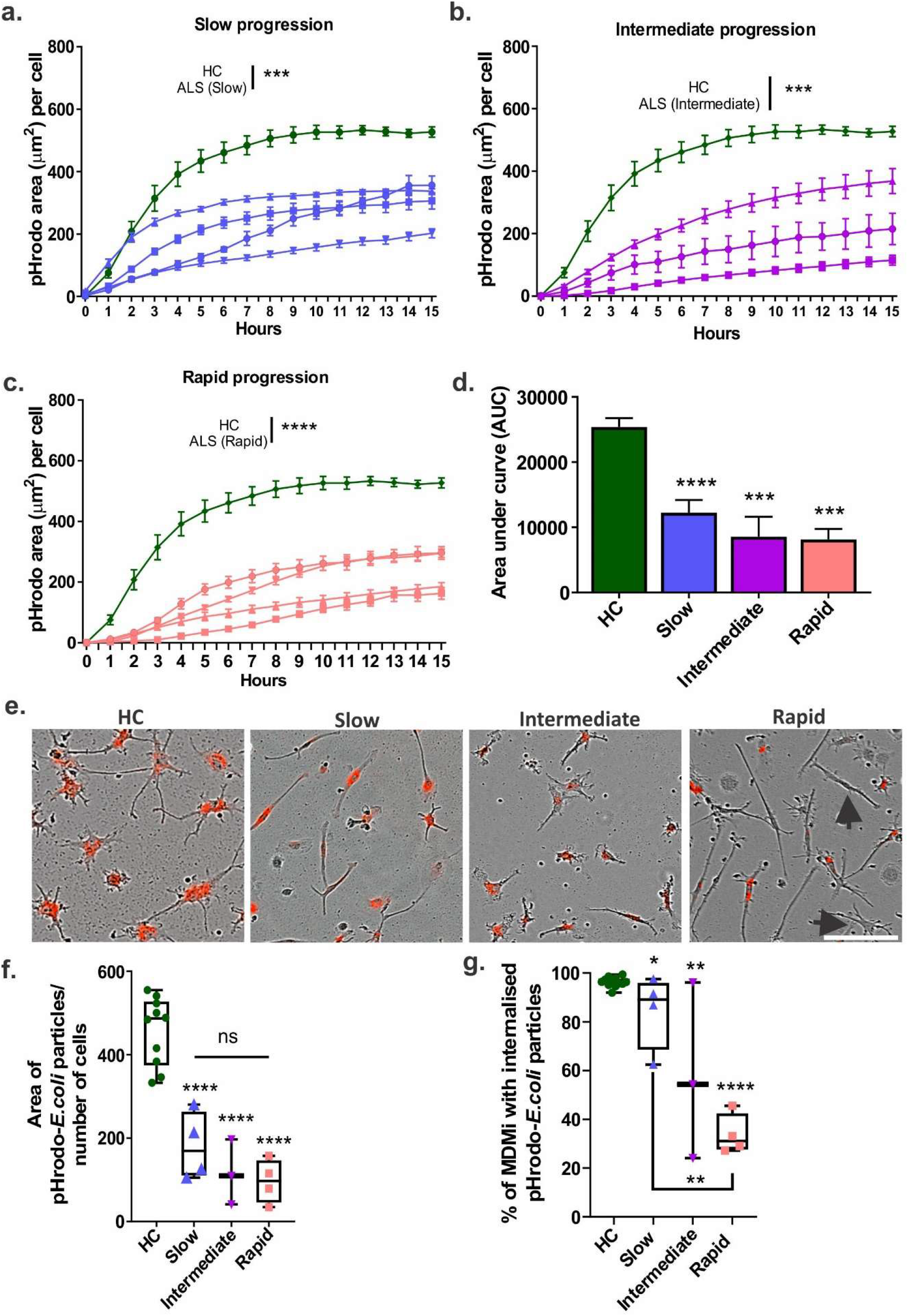
Impaired phagocytosis in ALS MDMi. Quantification of pHrodo-labelled *E.coli* particles phagocytosis over 15 hrs using live imaging in HC: n=10 (green) and **(a)** ALS slow: n=4 (blue), **(b)** intermediate: n=3 (purple), **(c)** rapid: n=4 (orange). **(d)** Uptake of pHrodo-labelled *E.coli* particles in HC and ALS disease subgroups MDMi was quantified using area under the curve. **(e)** Representative images of pHrodo-labelled *E.coli* particles (red) uptake by HC and ALS subgroup MDMi. Arrows show MDMi cells that have no uptake of pHrodo-labelled *E.coli* particles. **(f)** Area of pHrodo-labelled *E.coli* particles normalised against cell number, quantified using IncuCyte ZOOM in-built software. **(g)** Percentage of cells that phagocytose particles in HC and ALS MDMi. n=200 cells per individual. Statistical analysis between two groups was performed using Student’s *t* test and between multiple groups using one-way ANOVA. Values are mean ± SEM (*p < 0.05, **p < 0.01, ***p < 0.001, ****p < 0.0001). Scale bar= 50μm.

## References

Amlie-Wolf, A., Ryvkin, P., Tong, R., Dragomir, I., Suh, E., Xu, Y., … Lee, E. B. (2015). Transcriptomic Changes Due to Cytoplasmic TDP-43 Expression Reveal Dysregulation of Histone Transcripts and Nuclear Chromatin. PLOS ONE, 10(10), e0141836. doi:10.1371/journal.pone.0141836

Beers, D. R., & Appel, S. H. (2019). Immune dysregulation in amyotrophic lateral sclerosis: mechanisms and emerging therapies. The Lancet Neurology, 18(2), 211–220. doi:https://doi.org/10.1016/S1474-4422(18)30394-6

Bennett, M. L., Bennett, F. C., Liddelow, S. A., Ajami, B., Zamanian, J. L., Fernhoff, N. B., … Barres, B. A. (2016). New tools for studying microglia in the mouse and human CNS. Proceedings of the National Academy of Sciences, 113(12), E1738–E1746. doi:10.1073/pnas.1525528113

Brettschneider, J., Arai, K., Del Tredici, K., Toledo, J. B., Robinson, J. L., Lee, E. B., … Trojanowski, J. Q. (2014). TDP-43 pathology and neuronal loss in amyotrophic lateral sclerosis spinal cord. Acta neuropathologica, 128(3), 423–437. doi:10.1007/s00401-014-1299-6

Brettschneider, J., Toledo, J. B., Van Deerlin, V. M., Elman, L., McCluskey, L., Lee, V. M. Y., & Trojanowski, J. Q. (2012). Microglial Activation Correlates with Disease Progression and Upper Motor Neuron Clinical Symptoms in Amyotrophic Lateral Sclerosis. PLOS ONE, 7(6), e39216. doi:10.1371/journal.pone.0039216

Butovsky, O., Jedrychowski, M. P., Moore, C. S., Cialic, R., Lanser, A. J., Gabriely, G., … Weiner, H. L. (2014). Identification of a unique TGF-β–dependent molecular and functional signature in microglia. Nature Neuroscience, 17(1), 131–143. doi:10.1038/nn.3599

Butovsky, O., Siddiqui, S., Gabriely, G., Lanser, A. J., Dake, B., Murugaiyan, G., … Weiner, H. L. (2012). Modulating inflammatory monocytes with a unique microRNA gene signature ameliorates murine ALS. J Clin Invest, 122(9), 3063–3087. doi:10.1172/jci62636

Cedarbaum, J. M., Stambler, N., Malta, E., Fuller, C., Hilt, D., Thurmond, B., & Nakanishi, A. (1999). The ALSFRS-R: a revised ALS functional rating scale that incorporates assessments of respiratory function. Journal of the Neurological Sciences, 169(1), 13–21. doi:https://doi.org/10.1016/S0022-510X(99)00210-5

Cereda, C., Baiocchi, C., Bongioanni, P., Cova, E., Guareschi, S., Metelli, M. R., … Ceroni, M. (2008). TNF and sTNFR1/2 plasma levels in ALS patients. Journal of Neuroimmunology, 194(1), 123–131. doi: https://doi.org/10.1016/j.jneuroim.2007.10.028

Correia, A. S., Patel, P., Dutta, K., & Julien, J.-P. (2015). Inflammation Induces TDP-43 Mislocalization and Aggregation. PLOS ONE, 10(10), e0140248–e0140248. doi:10.1371/journal.pone.0140248

De Marco, G., Lomartire, A., Calvo, A., Risso, A., De Luca, E., Mostert, M., … Chiò, A. (2017). Monocytes of patients with amyotrophic lateral sclerosis linked to gene mutations display altered TDP-43 subcellular distribution. Neuropathology and Applied Neurobiology, 43(2), 133–153. doi:10.1111/nan.12328

De Marco, G., Lupino, E., Calvo, A., Moglia, C., Buccinnà, B., Grifoni, S., … Chiò, A. (2011). Cytoplasmic accumulation of TDP-43 in circulating lymphomonocytes of ALS patients with and without TARDBP mutations. Acta neuropathologica, 121(5), 611–622. doi:10.1007/s00401-010-0786-7

Deora, V., Lee, J. D., Albornoz, E. A., McAlary, L., Jagaraj, C. J., Robertson, A. A. B., … Woodruff, T. M. (2020). The microglial NLRP3 inflammasome is activated by amyotrophic lateral sclerosis proteins. Glia, 68(2), 407–421. doi:10.1002/glia.23728

Du, Y., Zhao, W., Thonhoff, J. R., Wang, J., Wen, S., & Appel, S. H. (2020). Increased activation ability of monocytes from ALS patients. Experimental Neurology, 328, 113259. doi:https://doi.org/10.1016/j.expneurol.2020.113259

Ehrhart, J., Smith, A. J., Kuzmin-Nichols, N., Zesiewicz, T. A., Jahan, I., Shytle, R. D., … Garbuzova-Davis, S. (2015). Humoral factors in ALS patients during disease progression. J Neuroinflammation, 12, 127. doi:10.1186/s12974-015-0350-4

Endo, F., Komine, O., Fujimori-Tonou, N., Katsuno, M., Jin, S., Watanabe, S., … Yamanaka, K. (2015). Astrocyte-Derived TGF-β1 Accelerates Disease Progression in ALS Mice by Interfering with the Neuroprotective Functions of Microglia and T Cells. Cell Reports, 11(4), 592–604. doi:https://doi.org/10.1016/j.celrep.2015.03.053

Fernandes-Alnemri, T., Wu, J., Yu, J. W., Datta, P., Miller, B., Jankowski, W., … Alnemri, E. S. (2007). The pyroptosome: a supramolecular assembly of ASC dimers mediating inflammatory cell death via caspase-1 activation. Cell Death Differ, 14(9), 1590–1604. doi:10.1038/sj.cdd.4402194

Frank-Cannon, T. C., Alto, L. T., McAlpine, F. E., & Tansey, M. G. (2009). Does neuroinflammation fan the flame in neurodegenerative diseases? Molecular Neurodegeneration, 4(1), 47. doi:10.1186/1750-1326-4-47

Fukazawa, H., Tsukie, T., Higashida, K., Fujikura, M., & Ono, S. (2013). An immunohistochemical study of increased tumor necrosis factor-α in the skin of patients with amyotrophic lateral sclerosis. Journal of Clinical Neuroscience, 20(10), 1371–1376. doi:https://doi.org/10.1016/j.jocn.2012.11.007

Ginhoux, F., Greter, M., Leboeuf, M., Nandi, S., See, P., Gokhan, S., … Merad, M. (2010). Fate Mapping Analysis Reveals That Adult Microglia Derive from Primitive Macrophages. Science, 330(6005), 841–845. doi:10.1126/science.1194637

Gosselin, D., Skola, D., Coufal, N. G., Holtman, I. R., Schlachetzki, J. C. M., Sajti, E., … Glass, C. K. (2017). An environment-dependent transcriptional network specifies human microglia identity. Science, 356(6344), eaal3222. doi:10.1126/science.aal3222

Haukedal, H., & Freude, K. (2019). Implications of Microglia in Amyotrophic Lateral Sclerosis and Frontotemporal Dementia. Journal of Molecular Biology, 431(9), 1818–1829. doi:https://doi.org/10.1016/j.jmb.2019.02.004

Hawrot, J., Imhof, S., & Wainger, B. J. (2020). Modeling cell-autonomous motor neuron phenotypes in ALS using iPSCs. Neurobiology of Disease, 134, 104680. doi:https://doi.org/10.1016/j.nbd.2019.104680

Heneka, M. T., Kummer, M. P., & Latz, E. (2014). Innate immune activation in neurodegenerative disease. Nat Rev Immunol, 14(7), 463–477. doi:10.1038/nri3705

Hijdra, D., Vorselaars, A. D. M., Grutters, J. C., Claessen, A. M. E., & Rijkers, G. T. (2012). Differential expression of TNFR1 (CD120a) and TNFR2 (CD120b) on subpopulations of human monocytes. Journal of Inflammation, 9(1), 38. doi:10.1186/1476-9255-9-38

Hu, Y., Cao, C., Qin, X.-Y., Yu, Y., Yuan, J., Zhao, Y., & Cheng, Y. (2017). Increased peripheral blood inflammatory cytokine levels in amyotrophic lateral sclerosis: a metaanalysis study. Scientific Reports, 7(1), 9094–9094. doi:10.1038/s41598-017-09097-1

Johann, S., Heitzer, M., Kanagaratnam, M., Goswami, A., Rizo, T., Weis, J., … Beyer, C. (2015). NLRP3 inflammasome is expressed by astrocytes in the SOD1 mouse model of ALS and in human sporadic ALS patients. Glia, 63(12), 2260–2273. doi:10.1002/glia.22891

Kadhim, H., Deltenre, P., Martin, J. J., & Sébire, G. (2016). In-situ expression of Interleukin-18 and associated mediators in the human brain of sALS patients: Hypothesis for a role for immune-inflammatory mechanisms. Med Hypotheses, 86, 14–17. doi:10.1016/j.mehy.2015.11.022

Kierdorf, K., Erny, D., Goldmann, T., Sander, V., Schulz, C., Perdiguero, E. G., … Prinz, M. (2013). Microglia emerge from erythromyeloid precursors via Pu.1- and Irf8-dependent pathways. Nature Neuroscience, 16(3), 273–280. doi:10.1038/nn.3318

Kimura, F., Fujimura, C., Ishida, S., Nakajima, H., Furutama, D., Uehara, H., … Hanafusa, T. (2006). Progression rate of ALSFRS-R at time of diagnosis predicts survival time in ALS. Neurology, 66(2), 265–267. doi:10.1212/01.wnl.0000194316.91908.8a

Kollewe, K., Mauss, U., Krampfl, K., Petri, S., Dengler, R., & Mohammadi, B. (2008). ALSFRS-R score and its ratio: A useful predictor for ALS-progression. Journal of the Neurological Sciences, 275(1), 69–73. doi:10.1016/j.jns.2008.07.016

Konopka, A., Whelan, D. R., Jamali, M. S., Perri, E., Shahheydari, H., Toth, R. P., … Atkin, J. D. (2020). Impaired NHEJ repair in amyotrophic lateral sclerosis is associated with TDP-43 mutations. Molecular Neurodegeneration, 15(1), 51. doi:10.1186/s13024-020-00386-4

Kuhle, J., Lindberg, R. L. P., Regeniter, A., Mehling, M., Steck, A. J., Kappos, L., & Czaplinski, A. (2009). Increased levels of inflammatory chemokines in amyotrophic lateral sclerosis. European Journal of Neurology, 16(6), 771–774. doi:10.1111/j.1468-1331.2009.02560.x

Lupton, M. K., Robinson, G. A., Adam, R. J., Rose, S., Byrne, G. J., Salvado, O., … Breakspear, M. (2020). A prospective cohort study of prodromal Alzheimer’s disease: Prospective Imaging Study of Ageing: Genes, Brain and Behaviour (PISA). medRxiv, 2020.2005.2004.20091140. doi:10.1101/2020.05.04.20091140

Mackenzie, I. R., Bigio, E. H., Ince, P. G., Geser, F., Neumann, M., Cairns, N. J., … Trojanowski, J. Q. (2007). Pathological TDP-43 distinguishes sporadic amyotrophic lateral sclerosis from amyotrophic lateral sclerosis with SOD1 mutations. Ann Neurol, 61(5), 427–434. doi:10.1002/ana.21147

Martin, L. J., Liu, Z., Chen, K., Price, A. C., Pan, Y., Swaby, J. A., & Golden, W. C. (2007). Motor neuron degeneration in amyotrophic lateral sclerosis mutant superoxide dismutase-1 transgenic mice: Mechanisms of mitochondriopathy and cell death. Journal of Comparative Neurology, 500(1), 20–46. doi:10.1002/cne.21160

McCombe, P. A., & Henderson, R. D. (2010). Effects of gender in amyotrophic lateral sclerosis. Gender Medicine, 7(6), 557–570. doi:https://doi.org/10.1016/j.genm.2010.11.010

McGill, R. B., Steyn, F. J., Ngo, S. T., Thorpe, K. A., Heggie, S., Ruitenberg, M. J., … Woodruff, T. M. (2020). Monocytes and neutrophils are associated with clinical features in amyotrophic lateral sclerosis. Brain Communications, 2(1). doi:10.1093/braincomms/fcaa013

Meroni, M., Crippa, V., Cristofani, R., Rusmini, P., Cicardi, M. E., Messi, E., … Galbiati, M. (2019). Transforming growth factor beta 1 signaling is altered in the spinal cord and muscle of amyotrophic lateral sclerosis mice and patients. Neurobiology of Aging, 82, 48–59. doi:https://doi.org/10.1016/j.neurobiolaging.2019.07.001

Meyer, B. C., Voss, K.-O., Tobias, F., Jakob, B., Durante, M., & Taucher-Scholz, G. (2013). Clustered DNA damage induces pan-nuclear H2AX phosphorylation mediated by ATM and DNa PK. Nucleic Acids Research, 41, 6109–6118.

Miller, P. G., Bonn, M. B., Franklin, C. L., Ericsson, A. C., & McKarns, S. C. (2015). TNFR2 Deficiency Acts in Concert with Gut Microbiota To Precipitate Spontaneous Sex-Biased Central Nervous System Demyelinating Autoimmune Disease. The Journal of Immunology, 195(10), 4668–4684. doi:10.4049/jimmunol.1501664

Mitchell, R. M., Freeman, W. M., Randazzo, W. T., Stephens, H. E., Beard, J. L., Simmons, Z., & Connor, J. R. (2009). A CSF biomarker panel for identification of patients with amyotrophic lateral sclerosis. Neurology, 72(1), 14–19. doi:10.1212/01.wnl.0000333251.36681.a5

Mitra, J., Guerrero, E. N., Hegde, P. M., Liachko, N. F., Wang, H., Vasquez, V., … Hegde, M. L. (2019). Motor neuron disease-associated loss of nuclear TDP-43 is linked to DNA double-strand break repair defects. Proceedings of the National Academy of Sciences, 116(10), 4696. doi:10.1073/pnas.1818415116

Neumann, M., Kwong, L. K., Truax, A. C., Vanmassenhove, B., Kretzschmar, H. A., Van Deerlin, V. M., … Lee, V. M.-Y. (2007). TDP-43-Positive White Matter Pathology in Frontotemporal Lobar Degeneration With Ubiquitin-Positive Inclusions. Journal of Neuropathology & Experimental Neurology, 66(3), 177–183. doi:10.1097/01.jnen.0000248554.45456.58

Nimmerjahn, A., Kirchhoff, F., & Helmchen, F. (2005). Resting microglial cells are highly dynamic surveillants of brain parenchyma in vivo. Science, 308(5726), 1314–1318. doi:10.1126/science.1110647

Ormel, P. R., Böttcher, C., Gigase, F. A. J., Missall, R. D., van Zuiden, W., Fernández Zapata, M. C., … de Witte, L. D. (2020). A characterization of the molecular phenotype and inflammatory response of schizophrenia patient-derived microglia-like cells. Brain, Behavior, and Immunity. doi:https://doi.org/10.1016/j.bbi.2020.08.012

Paolicelli, R. C., Jawaid, A., Henstridge, C. M., Valeri, A., Merlini, M., Robinson, J. L., … Rajendran, L. (2017). TDP-43 Depletion in Microglia Promotes Amyloid Clearance but Also Induces Synapse Loss. Neuron, 95(2), 297–308.e296. doi:10.1016/j.neuron.2017.05.037

Philips, T., & Rothstein, J. D. (2015). Rodent Models of Amyotrophic Lateral Sclerosis. Current protocols in pharmacology, 69, 5.67.61-65.67.21. doi:10.1002/0471141755.ph0567s69

Poloni, M., Facchetti, D., Mai, R., Micheli, A., Agnoletti, L., Francolini, G., … Bachetti, T. (2000). Circulating levels of tumour necrosis factor-alpha and its soluble receptors are increased in the blood of patients with amyotrophic lateral sclerosis. Neurosci Lett, 287(3), 211–214. doi:10.1016/s0304-3940(00)01177-0

Quek, H., Luff, J., Cheung, K., Kozlov, S., Gatei, M., Lee, C. S., … Lavin, M. F. (2017). A rat model of ataxia-telangiectasia: evidence for a neurodegenerative phenotype. Human Molecular Genetics, 26(1), 109–123. doi:10.1093/hmg/ddw371

Riancho, J., Castanedo-Vázquez, D., Gil-Bea, F., Tapia, O., Arozamena, J., Durán-Vían, C., … Lafarga, M. (2020). ALS-derived fibroblasts exhibit reduced proliferation rate, cytoplasmic TDP-43 aggregation and a higher susceptibility to DNA damage. J Neurol, 267(5), 1291–1299. doi:10.1007/s00415-020-09704-8

Ryan, K. J., White, C. C., Patel, K., Xu, J., Olah, M., Replogle, J. M., … Bradshaw, E. M. (2017). A human microglia-like cellular model for assessing the effects of neurodegenerative disease gene variants. Science Translational Medicine, 9(421), eaai7635. doi:10.1126/scitranslmed.aai7635

Schroder, K., & Tschopp, J. (2010). The Inflammasomes. Cell, 140(6), 821–832. doi:10.1016/j.cell.2010.01.040

Sellgren, C. M., Sheridan, S. D., Gracias, J., Xuan, D., Fu, T., & Perlis, R. H. (2017). Patientspecific models of microglia-mediated engulfment of synapses and neural progenitors. Molecular Psychiatry, 22(2), 170–177. doi:10.1038/mp.2016.220

Smethurst, P., Newcombe, J., Troakes, C., Simone, R., Chen, Y.-R., Patani, R., & Sidle, K. (2016). In vitro prion-like behaviour of TDP-43 in ALS. Neurobiology of Disease, 96, 236–247. doi:https://doi.org/10.1016/j.nbd.2016.08.007

Spiller, K. J., Restrepo, C. R., Khan, T., Dominique, M. A., Fang, T. C., Canter, R. G., … Lee, V. M. Y. (2018). Microglia-mediated recovery from ALS-relevant motor neuron degeneration in a mouse model of TDP-43 proteinopathy. Nature Neuroscience, 21(3), 329–340. doi:10.1038/s41593-018-0083-7

Svahn, A. J., Don, E. K., Badrock, A. P., Cole, N. J., Graeber, M. B., Yerbury, J. J., … Morsch, M. (2018). Nucleo-cytoplasmic transport of TDP-43 studied in real time: impaired microglia function leads to axonal spreading of TDP-43 in degenerating motor neurons. Acta neuropathologica, 136(3), 445–459. doi:10.1007/s00401-018-1875-2

Tortarolo, M., Lo Coco, D., Veglianese, P., Vallarola, A., Giordana, M. T., Marcon, G., … Bendotti, C. (2017). Amyotrophic Lateral Sclerosis, a Multisystem Pathology: Insights into the Role of TNF<i>α</i>. Mediators of Inflammation, 2017, 2985051. doi:10.1155/2017/2985051

Turner, M. R., Cagnin, A., Turkheimer, F. E., Miller, C. C. J., Shaw, C. E., Brooks, D. J., … Banati, R. B. (2004). Evidence of widespread cerebral microglial activation in amyotrophic lateral sclerosis: an [11C](R)-PK11195 positron emission tomography study. Neurobiology of Disease, 15(3), 601–609. doi:https://doi.org/10.1016/j.nbd.2003.12.012

Wang, M., Crisostomo, P. R., Markel, T. A., Wang, Y., & Meldrum, D. R. (2008). Mechanisms of sex differences in TNFR2-mediated cardioprotection. Circulation, 118(14 Suppl), S38–S45. doi:10.1161/CIRCULATIONAHA.107.756890

Wehrspaun, C. C., Haerty, W., & Ponting, C. P. (2015). Microglia recapitulate a hematopoietic master regulator network in the aging human frontal cortex. Neurobiology of Aging, 36(8), 2443.e2449–2443.e2420. doi:10.1016/j.neurobiolaging.2015.04.008

Young, K., & Morrison, H. (2018). Quantifying Microglia Morphology from Photomicrographs of Immunohistochemistry Prepared Tissue Using ImageJ. J Vis Exp (136). doi:10.3791/57648

Zhao, W., Beers, D. R., Bell, S., Wang, J., Wen, S., Baloh, R. H., & Appel, S. H. (2015). TDP-43 activates microglia through NF-κB and NLRP3 inflammasome. Exp Neurol, 273, 24–35. doi:10.1016/j.expneurol.2015.07.019

Zhao, W., Beers, D. R., Hooten, K. G., Sieglaff, D. H., Zhang, A., Kalyana-Sundaram, S., … Appel, S. H. (2017). Characterization of Gene Expression Phenotype in Amyotrophic Lateral Sclerosis Monocytes. JAMA Neurol, 74(6), 677–685. doi:10.1001/jamaneurol.2017.0357

Zhu, J., Cynader, M. S., & Jia, W. (2015). TDP-43 Inhibits NF-κB Activity by Blocking p65 Nuclear Translocation. PLOS ONE, 10(11), e0142296. doi:10.1371/journal.pone.0142296

Zondler, L., Müller, K., Khalaji, S., Bliederhäuser, C., Ruf, W. P., Grozdanov, V., … Weishaupt, J. H. (2016). Peripheral monocytes are functionally altered and invade the CNS in ALS patients. Acta neuropathologica, 132(3), 391–411. doi:10.1007/s00401-016-1548-y

Zusso, M., Methot, L., Lo, R., Greenhalgh, A. D., David, S., & Stifani, S. (2012). Regulation of postnatal forebrain amoeboid microglial cell proliferation and development by the transcription factor Runx1. The Journal of neuroscience: the official journal of the Society for Neuroscience, 32(33), 11285–11298. doi:10.1523/JNEUROSCI.6182-11.2012

